# Inferring protein sequence-function relationships with large-scale positive-unlabeled learning

**DOI:** 10.1101/2020.08.19.257642

**Authors:** Hyebin Song, Bennett J. Bremer, Emily C. Hinds, Garvesh Raskutti, Philip A. Romero

## Abstract

Machine learning can infer how protein sequence maps to function without requiring a detailed understanding of the underlying physical or biological mechanisms. It’s challenging to apply existing supervised learning frameworks to large-scale experimental data generated by deep mutational scanning (DMS) and related methods. DMS data often contain high dimensional and correlated sequence variables, experimental sampling error and bias, and the presence of missing data. Importantly, most DMS data do not contain examples of negative sequences, making it challenging to directly estimate how sequence affects function. Here, we develop a positive-unlabeled (PU) learning framework to infer sequence-function relationships from large-scale DMS data. Our PU learning method displays excellent predictive performance across ten large-scale sequence-function data sets, representing proteins of different folds, functions, and library types. The estimated parameters pinpoint key residues that dictate protein structure and function. Finally, we apply our statistical sequence-function model to design highly stabilized enzymes.

## Introduction

A protein’s sequence of amino acids encodes its function. This “function” could refer to a protein’s natural biological function, or it could also be any other property including binding affinity toward a particular ligand, thermodynamic stability, or catalytic activity. A detailed understanding of how these functions are encoded would allow us to more accurately reconstruct the tree of life, diagnose genetic diseases before they manifest symptoms, and design new proteins with useful properties. The mapping from protein sequence to function is extraordinarily complex because it involves thousands of molecular interactions that are dynamically coupled across multiple length and time scales.

Machine learning can infer how protein sequence encodes function without needing to understand the underlying biophysical mechanisms (Yang et al. 2019, Mazurenko et al. 2020). These learning methods can be broadly categorized into unsupervised and supervised depending on whether the data points are labeled. In the protein context, unsupervised methods learn from examples of sequences that share some common function/property, while supervised methods learn from sequence-function examples. Unsupervised methods are often trained on natural sequence data derived from large genomic databases, and effectively learn the rules of folding/function for a given protein family (Morcos et al. 2011, Hopf et al. 2017, Riesselman et al. 2018). In contrast, supervised methods are trained directly on sequence-function examples, and therefore, can learn the mapping to a particular protein property or set of properties. This capability to predict a target protein property is important in protein engineering, which seeks to design and optimize non-natural protein functions. Supervised models have been used to rationally engineer proteinases with improved activity at elevated temperatures, cytochrome P450s with enhanced stability, carbonic anhydrases for industrial carbon capture, and bacteriorhodopsins for optogenetics (Liao et al. 2007, Romero et al. 2013, Alvizo et al. 2014, Bedbrook et al. 2019).

The accuracy and resolution of statistical models improve with increasing data; however, existing supervised methods cannot learn from large-scale sequence-function data generated by deep mutational scanning (DMS) and related methods. DMS combines high-throughput screening and next-generating DNA sequencing to experimentally map sequence-function relationships for thousands to millions of protein variants (Fowler & Fields 2014, Boucher et al. 2014, Weile & Roth 2018). In principle, DMS data should provide rich sequence-function information for training supervised models. However, learning from DMS data is challenging due to its scale and dimensionality, correlations between sequence variables, sampling error caused by low numbers of observations, and missing/low quality sequence information. In addition, most DMS data sets do not contain negative sequence examples because these sequences are difficult or impossible to obtain using high-throughput screening/selection methods. These negative sequences are important to directly infer how sequence maps to function. Hence DMS data are neither amenable to supervised learning due to the lack of negative sequences, nor unsupervised learning since many sequences have positive labels.

In this paper, we present a novel supervised learning framework for inferring sequence-function relationships from large-scale data generated by deep mutational scanning (DMS). We categorize DMS data as *positive-unlabeled* (PU) data because it contains examples of both positive sequences and sequences without labels. Learning from PU data has applications in domains such as text mining, gene identification, marketing, and ecological modeling (Liu et al. 2003, Mordelet & Vert 2011, Yi et al. 2017, Ward et al. 2009). We develop a PU learning method that models a protein’s function as an unobserved latent variable, and then infers how sequence maps to this latent function by maximizing an observed likelihood function. Our learned PU models displayed excellent predictive ability and stability across ten diverse DMS data sets. The PU model’s parameters describe how amino acid substitutions affect protein function, and the significance of these parameter estimates can be evaluated using statistical hypothesis testing. We demonstrate the extrapolative power of the learned sequence-function mapping by designing enzymes with significantly increased thermostability.

## Results

### A statistical framework for learning sequence-function relationships

Supervised learning methods can infer how sequence maps to function from a set of experimental sequence-function examples. However, it’s challenging to apply existing learning methods to large-scale data generated by deep mutational scanning (DMS) due to the lack of negative sequence examples. We use the term DMS to broadly refer to any experiment that maps sequence-function relationships using a combination of gene mutagenesis, high-throughput screening/selections, and next-generation DNA sequencing. An overview of a standard DMS experiment is illustrated in Figure 1a. This section describes the DMS data generation process, introduces key statistical variables, and proposes a positive-unlabeled approach to learn from DMS data.

**Figure 1:**
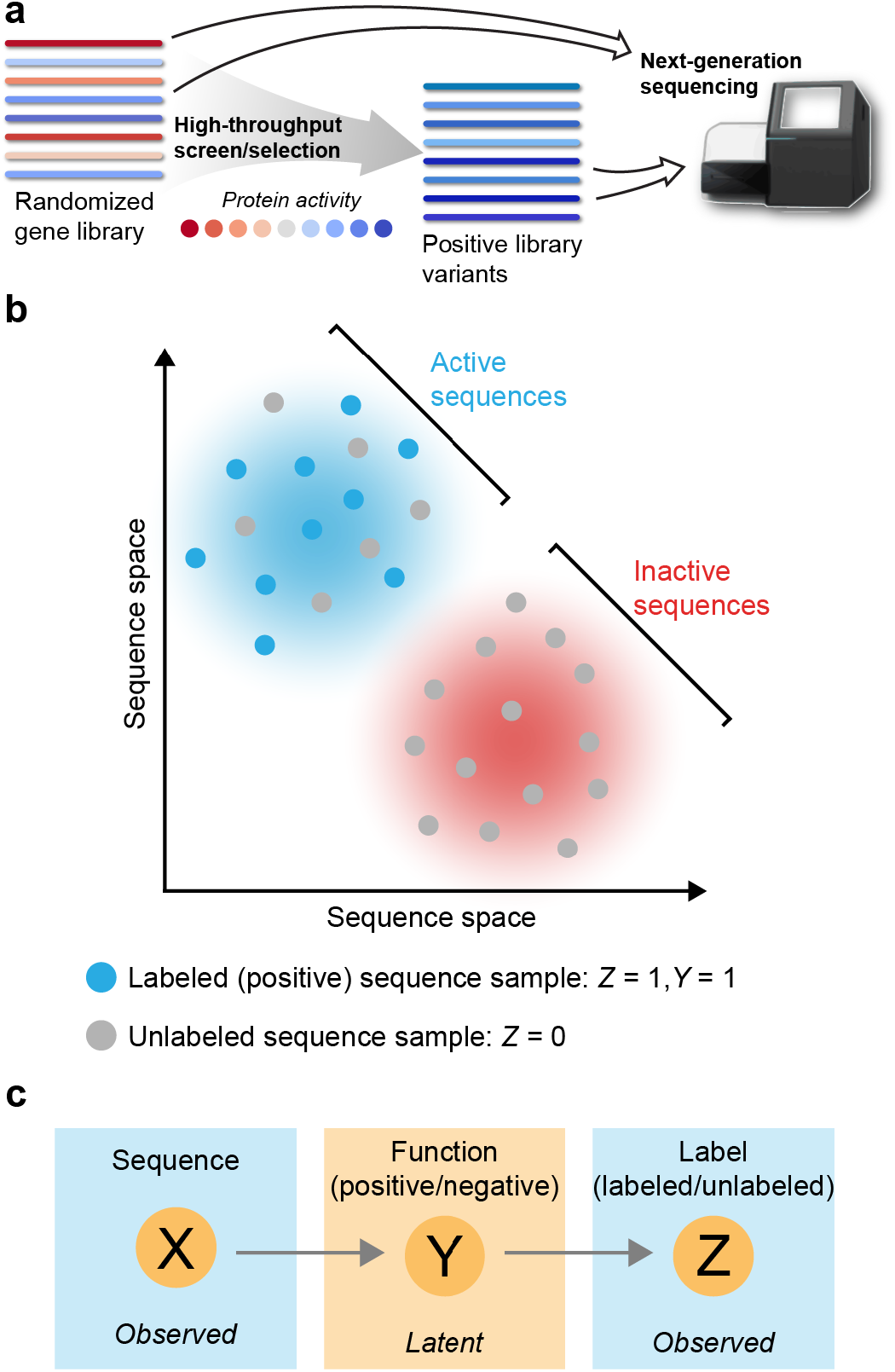
(a) Overview of a typical deep mutational scanning (DMS) experiment. DMS experiments start with a large library of gene variants that display a range of activities. The gene library is then expressed and passed through a high-throughput screen or selection that isolates the positive variants. The activity threshold to be categorized as positive will depend on the details of the particular high-throughput screen/selection. Importantly, it is often difficult or impossible to experimentally isolate negative sequences. Genes from the initial library and the isolated positive variants are then extracted and analyzed using next-generation DNA sequencing. DMS experiments generate thousands to millions of sequence examples from both the initial and positive sets. (b) DMS experiments sample sequences from protein sequence space. The resulting data contain positive labeled sequence examples (*Y* = 1, *Z* = 1), and unlabeled sequence examples (*Z* = 0) that contain a mixture of active and inactive sequences. (c) The relationships between variables representing protein sequences (*X*), latent function (*Y*), and the observed labels (*Z*). *Y* is not directly observed in DMS experiments and must be inferred from *X* and *Z*.

A protein’s biochemical activity *A*_*i*_ is a function of its amino acid sequence, i.e. *A*_*i*_ = *f*(*x*_*i*_), where *x*_*i*_ is a vector that specifies a protein’s amino acid sequence (see Materials and Methods). Now suppose a protein sequence can be categorized as active or inactive depending on whether its activity *A*_*i*_ falls above/below a defined activity threshold *t*. There is also some error in experimentally determining whether a sequence is active or inactive. We define a protein’s experimentally measured functional response as:

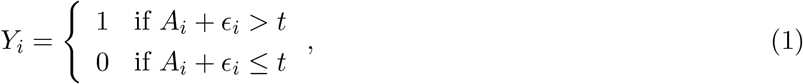

where *ϵ*_*i*_ is the random error associated with experimentally characterizing the *i*th sequence. We say a sequence is “positive” if its experimentally measured activity exceeds the threshold *t* (i.e. *Y*_*i*_ = 1) and “negative” otherwise. Note that we make subtle distinction between the terms active/inactive and positive/negative: we use active/inactive to describe the true functional state of a protein and positive/negative to indicate the result of an experimental measurement.

A DMS experiment starts with an initial library of sequences that each maps to a particular activity value and can be categorized as active or inactive. We refer to this initial library as “unlabeled” because it contains an unknown mixture of active and inactive sequences. A high-throughput screen/selection then samples this initial unlabeled library to obtain examples of positive sequences. Importantly, it is often difficult or impossible to isolate negative sequences because most experimental methods are designed to identify positive sequences (e.g. growth selections). We refer to the sampled positive sequences as “labeled” because they are known to have positive labels. The sequences within the initial unlabeled set and positive labeled set are then determined using next-generation sequencing. The final data contains *n*_*u*_ sequences sampled from the unlabeled set and *n*_*p*_ sequences sampled from the positive labeled set (Figure 1b).

We aim to learn from DMS data to understand how amino acid sequence maps to function (i.e. infer *f*). If the data consisted of (*X*_*i*_, *Y*_*i*_) pairs, we could simply train a binary classifier such as a logistic regression model or a multilayer perceptron. However, DMS experiments do not reveal the true functional response (*Y*), but instead only provide examples of positive sequences. The lack of negative sequence examples results in a mis-specified binary classification problem and makes it challenging to directly infer how sequence maps to function.

We propose a positive-unlabeled approach to learn from DMS data. Positive-unlabeled (PU) learning estimates how input variables affect the positive-negative response from positive and unlabeled data (Liu et al. 2003, Elkan & Noto 2008, Song & Raskutti 2018). We introduce a new binary variable *Z* that specifies whether a sequence is labeled (*Z* = 1) or unlabeled (*Z* = 0). DMS experiments effectively generate (*X*_*i*_, *Z*_*i*_) pairs with an unobserved functional response *Y*_*i*_ (Figure 1c). From the set-up of the problem, *Z* and *Y* are closely related. In particular, all labeled sequences are positive, and the proportion of positive sequences in the unlabeled set is the same as that of the initial library. In other words,

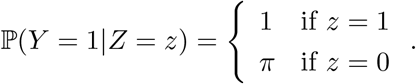

where *π* is the proportion of positive sequences in the initial library. We aim to infer *f* using the observed examples 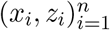, where *n* = *n*_*p*_ + *n*_*u*_ denotes the total number of sequence examples.

### Novel algorithms for large-scale positive-unlabeled learning

Sequence-function data obtained by deep mutational scanning (DMS) typically contains sequence examples from the initial (unlabeled) library and positive sequences (Figure 1). We develop novel algorithms for learning the sequence-function mapping from this positive-unlabeled sequence data. Our approach utilizes the distributional relationship between *X*, *Y*, and *Z* to infer the latent functional response *y*_*i*_ from the observed labels 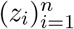 and amino acid sequences 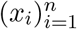, and then infer the *f* that best describes the latent responses 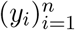.

We model *f* as a linear function of amino acid sequence 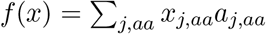, and use a logistic function to describe the probability that a sequence is positive:

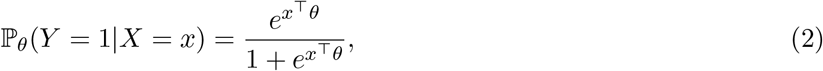

where 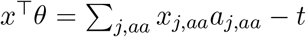 represents a relative activity level of amino acid sequence *x* with respect to the activity threshold *t* in (1), and *θ* parameterizes the effect of making an amino acid substitution from a defined reference sequence (see Materials and Methods). The model’s parameters (*θ*) are closely related to the site-wise enrichment scores that are commonly used to analyze DMS data (Bloom 2015, Klesmith & Hackel 2019, Wrenbeck et al. 2017, Abriata et al. 2016). However, site-wise enrichment is a biased estimator for *θ* because it is derived from positive-unlabeled data and makes strong assumptions about the independence between sequence positions. We derive the mathematical relationship between site-wise enrichment and a mutation’s true effect (*θ*) in the STAR Methods.

We take a likelihood-based approach for estimating *θ* from the observed examples 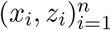. To account for the fact that the true responses 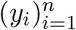 are latent, we use an observed likelihood that is a product of the marginalized probabilities (Ward et al. 2009)

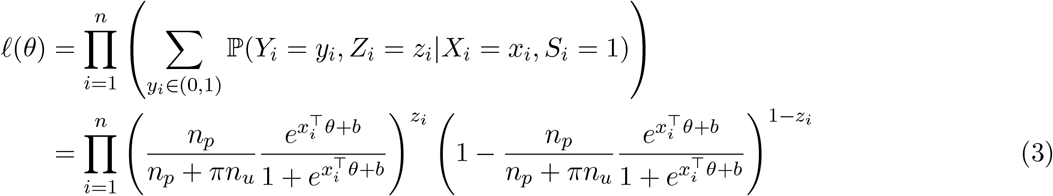

for 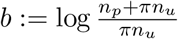, where *S*_*i*_ ∈ (0, 1) is an indicator variable representing whether the ith example is present in the data. We use the maximum likelihood approach to estimate *θ*. In particular, we minimize the *negative* observed log-likelihood and define the estimated coefficients 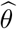 as

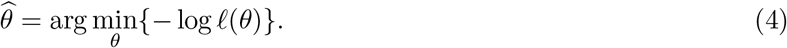

The negative observed log-likelihood, 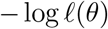, is a non-convex function of *θ*. Obtaining a global minimizer of a non-convex function is in general a challenging problem, so the feasibility of obtaining 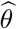 is not immediate. We previously found similar classes of problems can be solved when the likelihood function is calculated with sufficiently large sample size *n* (Song et al. 2019). In these cases, any stationary point is the global minimizer with high probability. Since our sequence-function data sets typically contain millions of observations, we can find the maximum likelihood estimate by identifying a stationary point of the negative observed log-likelihood.

We solved this optimization problem using the Majorization-Minimization (MM) algorithm to obtain a stationary point of the negative observed log-likelihood function (Eqn 3). Note that the likelihood (Eqn 3) involves the hyperparameter *π*, the proportion of positive sequences in the unlabeled set. *π* was experimentally determined for some data sets, and we used this value for the hyperparameter if it was available. Otherwise, we carried out a grid search over *π* values and chose the *π* value that maximized the area under the receiver operating characteristic curve. We used the learned model parameters to calculate p-values to test whether amino acid substitutions have a significant impact on protein function. These p-values were adjusted using the Benjamini-Hochberg (BH) procedure to account for multiple hypothesis testing (Materials and Methods). An overview of our data processing, parameter estimation, and model analysis workflow is provided in Supplementary Figure 2.

### Relationship between learned PU model parameters and site-wise enrichment scores

There is a close connection between our PU model’s parameters and the site-wise enrichment scores that are commonly used to analyze deep mutational scanning (DMS) data (Bloom 2015, Klesmith & Hackel 2019, Wrenbeck et al. 2017, Abriata et al. 2016). Both quantities evaluate how amino acid substitutions affect a protein’s functional response (i.e. estimate 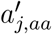). However, our PU model provides a consistent estimate of 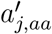 because it directly models the true positive-negative (PN) response and considers the conditional effects of amino acid substitutions. In this section, we define site-wise enrichment scores, contrast them with the PU model parameters, and identify two different sources of bias in their estimate of 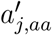.

Site-wise enrichment scores are calculated using marginal amino acid frequencies:

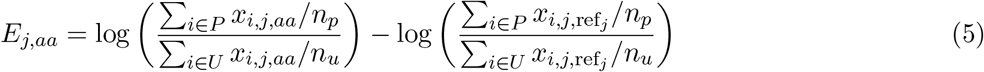

where *P* and *U* are the positive and unlabeled sets of sequences, respectively. *E*_*j,aa*_ compares the prevalence of amino acid *aa* with the reference sequence (typically wild-type) in the positive and unlabeled sets. A residue with a negative enrichment score is underrepresented in the positive set, and therefore associated with decreased protein activity. Conversely, a residue with a positive enrichment score is associated with the increased protein activity. These enrichment scores provide a simple and convenient method to compare frequencies before/after selection and estimate the effects of amino acid substitutions.

Enrichment and our PU learning method capture different response variables related to protein function (Figure 2a). Enrichment models the positive-unlabeled (PU) response, and since all labeled sequences are positive, this is equivalent to modeling a sequence’s label *Z*. In contrast, our PU learning method directly models the positive-negative (PN) response by inferring the latent function *Y* from observed labels *Z*. The presence of positive sequences in the unlabeled set causes enrichment-based methods to provide attenuated estimates of an amino acid substition’s effect (see STAR Methods). This leads to a decision boundary that is shifted toward the positive class and results in misclassified sequences. Our PU learning method models the true PN response, and thus provides an unbiased estimate of a substitution’s effect 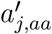.

**Figure 2:**
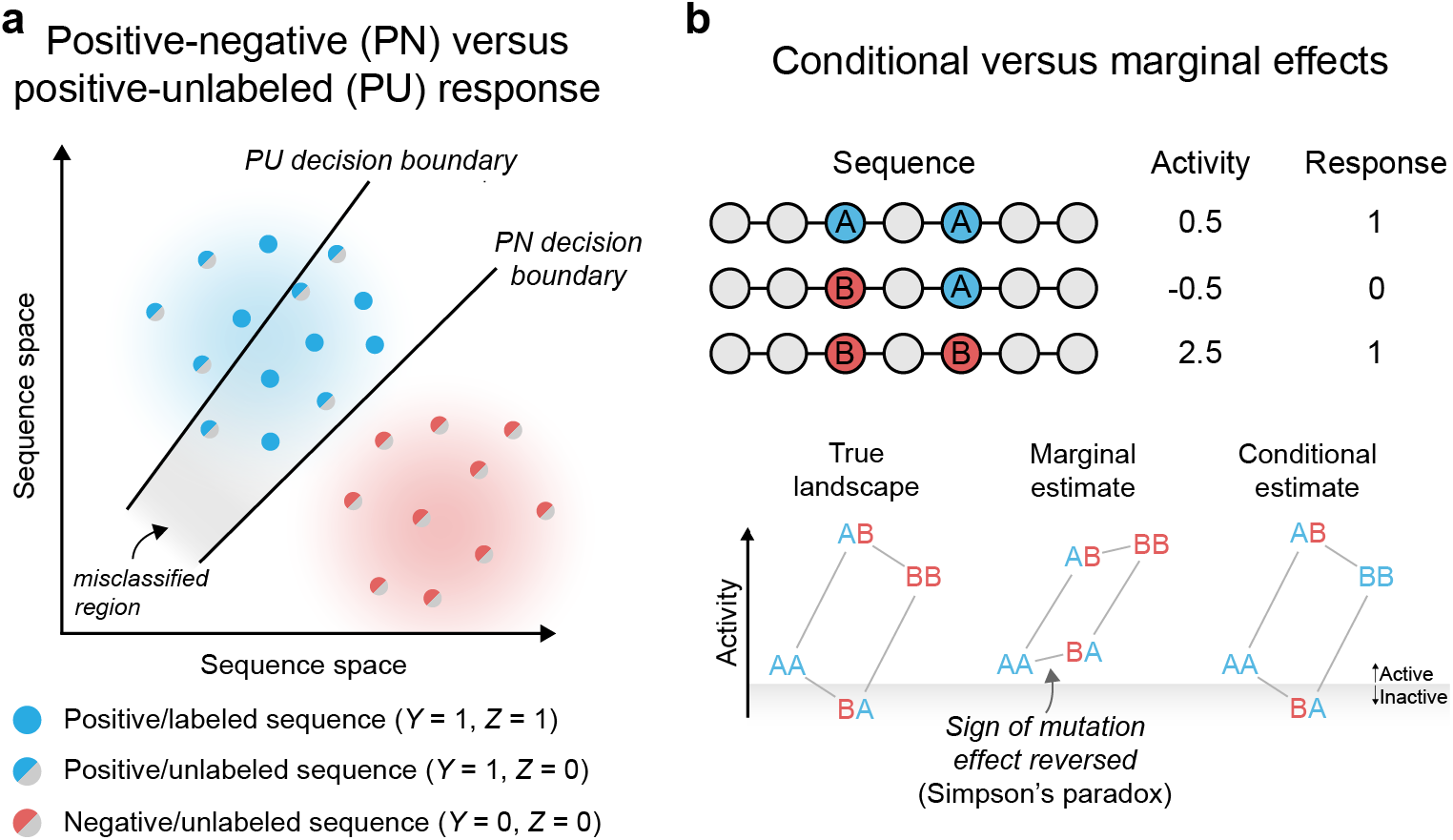
Contrasting PU learning versus enrichment-based estimates of protein function. (a) PU learning models the true positive-negative (PN) response, while enrichment-based estimates capture the positive-unlabeled (PU) response. Modeling the PU response gives rise to a decision boundary that is shifted toward the positive class, resulting in positive sequences that are misclassified as negative. (b) PU learning estimates the conditional effect of a mutation, while site-wise enrichment estimates the marginal effect. Marginal estimates are biased and in extreme cases can result in a sign reversal phenomenon known as Simpson’s paradox. In the example, we consider amino acid substitutions A→B at two independent sites in a protein. If we observe sequences AA, BA, and BB, the marginal estimate will reverse the sign of substitution A→B at the first position. The marginal model will also misclassify sequence BA as positive, even though it was observed to be negative. In contrast, the conditional estimate correctly models the true protein function landscape.

The second key difference between PU model parameters and enrichment is related to marginal versus conditional effects. Our PU learning method estimates the effect of an amino acid substitution from a defined sequence background, typically wild-type. This is a *conditional* estimate because the effect of the mutation is conditioned on all other sites in the protein. This conditional effect provides an unbiased estimate of an amino acid substitution’s effect 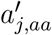. In contrast, site-wise enrichment estimates the effect of an amino acid substitution in combination with averaged effects from all other sites in the data set. These *marginal* estimates include the true effect 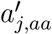, in addition to indirect effects from other sequence positions. These indirect effects lead to bias in the estimate of 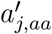 (see STAR Methods). In more extreme instances, marginal estimates can reverse the sign of an effect and lead to incorrect conclusions about whether a substitution is beneficial or deleterious (Figure 2b). This effect reversal is referred to as Simpson’s paradox.

### Learning from large-scale sequence-function data

We applied our PU learning method to infer the sequence-activity mapping from ten large sequence-function data sets (see Supplementary Table 1). These data sets represent proteins of diverse folds/functions, were generated using different library mutagenesis methods, span several orders of magnitude in size, and have varying levels of missing sequence information. The PU models displayed excellent predictive ability on all ten data sets, with cross-validated area under the receiver operating characteristic curve (ROC-AUC) ranging from 0.68 to 0.98 (Figure 3ab). For comparison, we also evaluated predictions from structure-based (Alford et al. 2017) and unsupervised learning methods (Hopf et al. 2017, Riesselman et al. 2018). Rosetta, EVmutation, and DeepSequence all displayed substantially lower AUC values than the PU model (Figure 3b, Supplementary Figure 3a).

**Figure 3:**
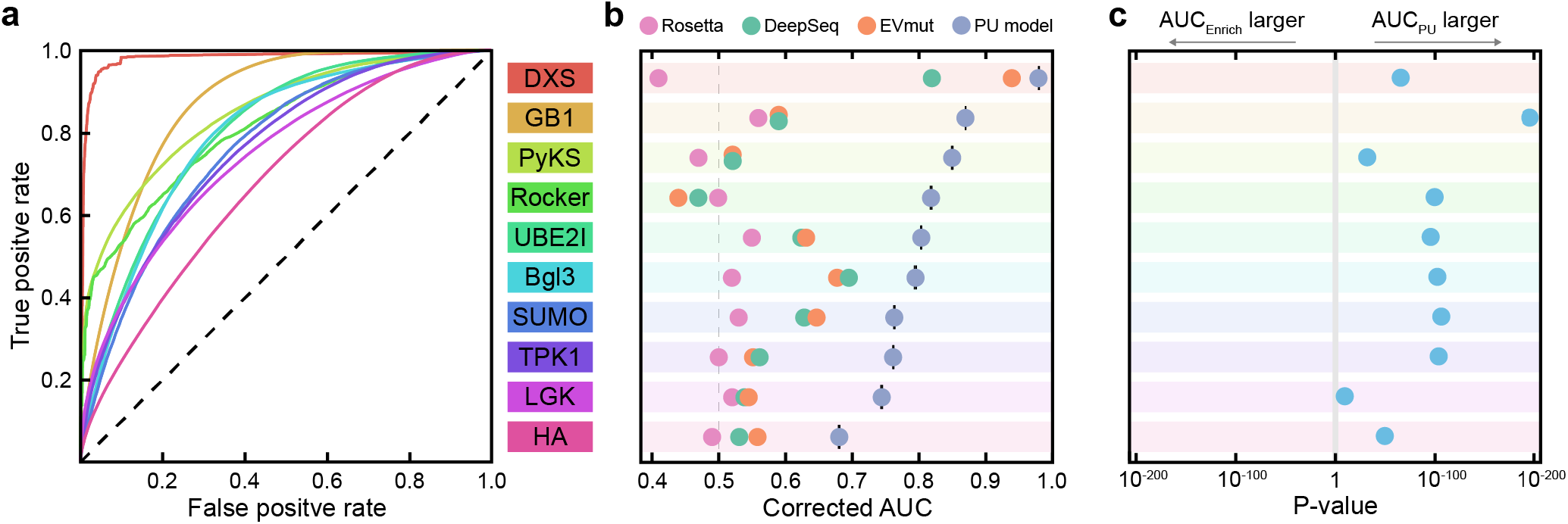
Performance of the PU learning method across protein data sets. (a) Receiver operating characteristic (ROC) curves for the ten tested protein data sets. ROC curves were generated using 10-fold cross-validation and corrected to account for PU data (See Materials and Methods and Supplementary Figure 1). (b) The PU model’s corrected ROC-AUC values range from 0.68 to 0.98, and outperform structure-based (Rosetta) and unsupervised learning methods (EVmutation and DeepSequence). (c) A statistical comparison between PU model predictions and site-wise enrichment. The PU model outperformed enrichment on all ten tested data sets, with *p*< 10^−9^.

Our PU learning method estimates how mutations affect a protein’s functional response. This PU estimate is closely related to the site-wise enrichment scores that are commonly used to analyze DMS data. We compared the predictive ability of the PU model versus enrichment using a corrected cross-validation test. We found the PU model predictions were better than enrichment for all ten data sets, with *p*< 10^−9^ (Figure 3c). However, the PU models’ AUCs were only marginally higher than enrichment, with AUC differences ranging from 0.002 to 0.017 (Supplementary Figure 3b).

We evaluated the robustness of the learned PU models to data sampling and the hyperparameter *π*. We analyzed the stability of each model’s parameter estimates by calculating the coefficient of variation (CV) across different cross-validation folds for all significant parameters (i.e. BH-adjusted *p* < 0.05). The parameters displayed average absolute CVs ranging from 0.01 to 0.08 (Supplementary Figure 3c), indicating the estimates were highly insensitive to different training sets. We also evaluated the feature selection stability by computing the average fraction of commonly selected features across different cross-validation folds (Materials and Methods). We found the selected features were nearly identical across cross-validation folds for each data set (Supplementary Figure 3d). Finally, we tested how the choice of the hyperparameter *π* (experimentally determined or estimated) affects the learned PU models. We found the parameter values estimated using our chosen *π* value were highly correlated with parameters estimated across the entire range of *π* values tested (Supplementary Figure 3e).

### Learned parameters relate to protein structure and function

The B1 domain of protein G (GB1) is a small 8 kDa alpha-beta roll that binds to IgG. We performed further analysis relating the learned GB1 model with protein G structure and function. The PU model’s coefficients describe how an amino acid substitution (mutation) affects the protein’s functional response (Eqn 7). A negative coefficient indicates that a substitution decreases protein activity, while a substitution with a positive coefficient increases activity. We found that most amino acid substitutions in GB1 are slightly deleterious, while a smaller subset is highly deleterious (Figure 4a). Each position in the amino acid sequence displayed a range of mutational effects (Figure 4b). Substitutions to proline are the most deleterious on average (presumably because they disrupt protein structure), followed by substitutions to the acidic amino acids. We found a site’s average mutational effect is highly dependent on its location in the three-dimensional structure. Sites with large average mutational effects tend to be located in either the protein core or the IgG binding interface (Figure 4bc).

**Figure 4:**
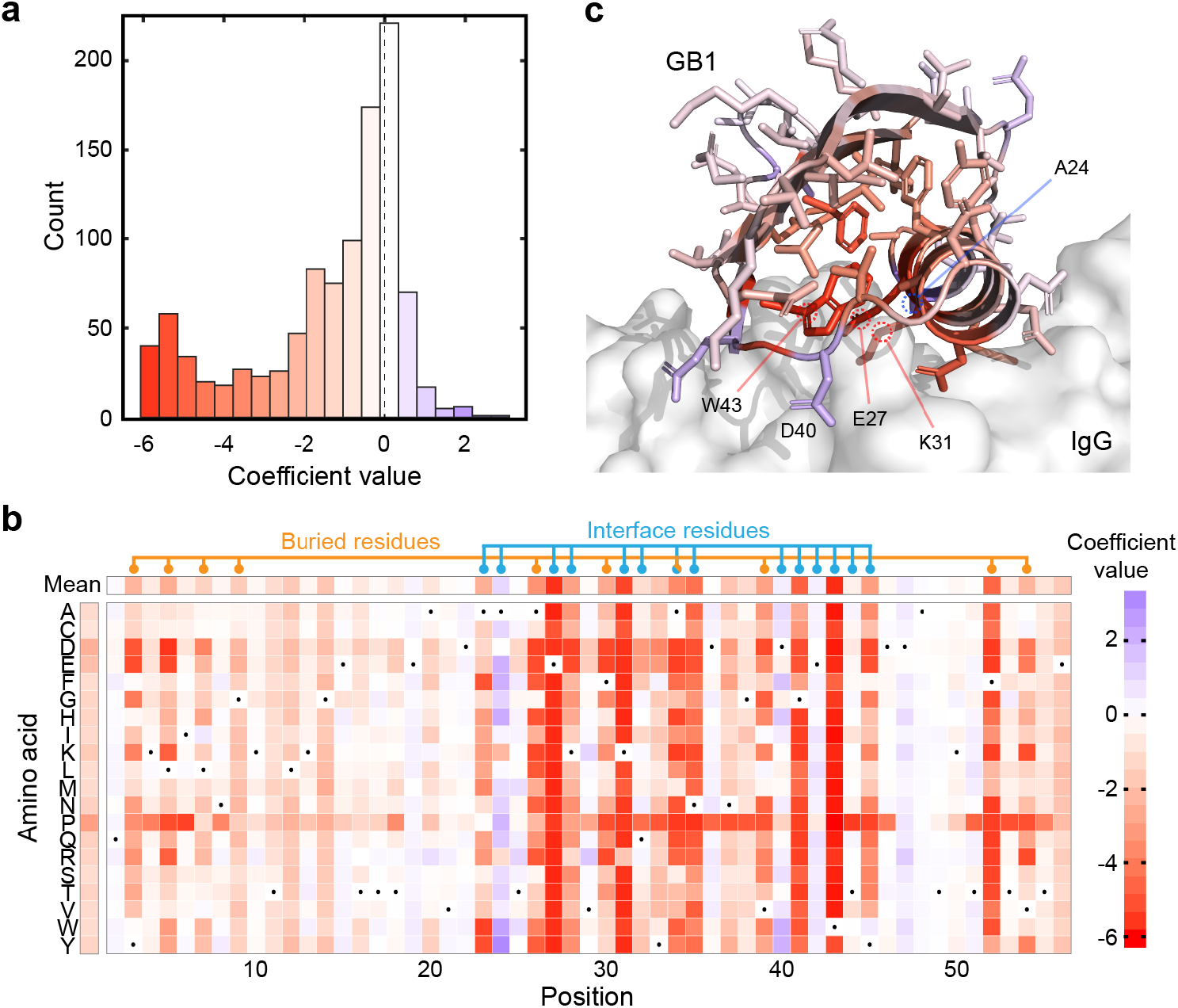
Model parameters relate to GB1 structure and function. (a) The distribution of model coefficients. Most coefficients have a relatively small magnitude, while a substantial fraction of coefficients have a large negative effect. (b) A heatmap of the GB1 model coefficients. The wild-type amino acid is depicted with a black dot. Buried and interface residues tend to have larger magnitude coefficients, indicating their important role in GB1 function. Buried and interface residues were determined from the protein G crystal structure (PDB ID: 1FCC). Buried residues were defined as having a relative solvent accessibility less than 0.1. Interface residues were defined as having a heavy atom within 4Å of IgG. (c) The site-wise average model coefficients mapped onto the protein G crystal structure (PDB ID: 1FCC). The IgG binding partner is depicted as a grey surface. Residues in the protein core and binding interface tend to have the largest average coefficients.

GB1 residues E27, K31, and W43 have the most negative average mutational effect, suggesting that many substitutions at these positions are highly deleterious. Consistent with the model results, these three residues are known to form key hydrogen bonds and salt bridges with the IgG ligand and make the largest contributions to the free energy of binding (Sauer-Eriksson et al. 1995, Sloan & Hellinga 1999). Residues A24 and D40 have the largest positive average mutational effect, with many substitutions that are predicted to increase GB1 activity. Both of these sites are located in the IgG-binding interface. Previous studies have identified residue position 24 to play a key role in IgG binding, and substitutions from A24 can increase binding affinity through improved ionic interactions (Sauer-Eriksson et al. 1995). Furthermore, high-affinity computationally designed protein G variants have substitutions at position 24 (Jha et al. 2014). The model parameters suggest that residue D40 prefers substitution to aromatic amino acids (Figure 4b). Inspection of the crystal structure suggests these mutations could form potential interactions (pi-pi, cation-pi) with a nearby histidine in IgG (Supplementary Figure 4).

### Statistics-based protein design

Our PU learning method provides a quantitative description of the sequence-function mapping. Importantly, the model also captures statistical uncertainties arising from undersampling and correlated sequence variables. Here, we develop a protein design framework that leverages this statistical sequence-function information.

We trained a PU model on a Bgl3 deep mutational scan that had been performed at an elevated temperature (Bgl3-HT, Supplementary Table 1) (Romero et al. 2015). Under these experimental conditions, the positive class corresponds to Bgl3 sequences with a high thermal tolerance, and therefore the model should learn how amino acid substitutions affect thermostability. The learned PU model displayed excellent predictive ability (Corrected AUC of 0.72, Supplementary Figure 5a).

We applied the PU model to design Bgl3 variants based on either coefficient magnitudes or p-values (Figure 5a). The coefficient-based design (Bgl.cf) contained ten amino acid substitutions corresponding to the ten largest positive coefficients. The p-value-based design (Bgl.pv) contained ten substitutions corresponding to the ten positive coefficients with the smallest p-values. We also designed a sequence that contained the ten substitutions with the largest enrichment scores (Bgl.en). The Bgl.cf and Bgl.en designs contained six common substitutions, while the substitutions in Bgl.pv were distinct from the other two (Supplementary Figure 5b). The substitutions within these three designs are generally distributed throughout the protein structure (Figure 5b); however, there appears to be some bias for the coefficient/enrichment designs to choose substitutions in the termini.

**Figure 5:**
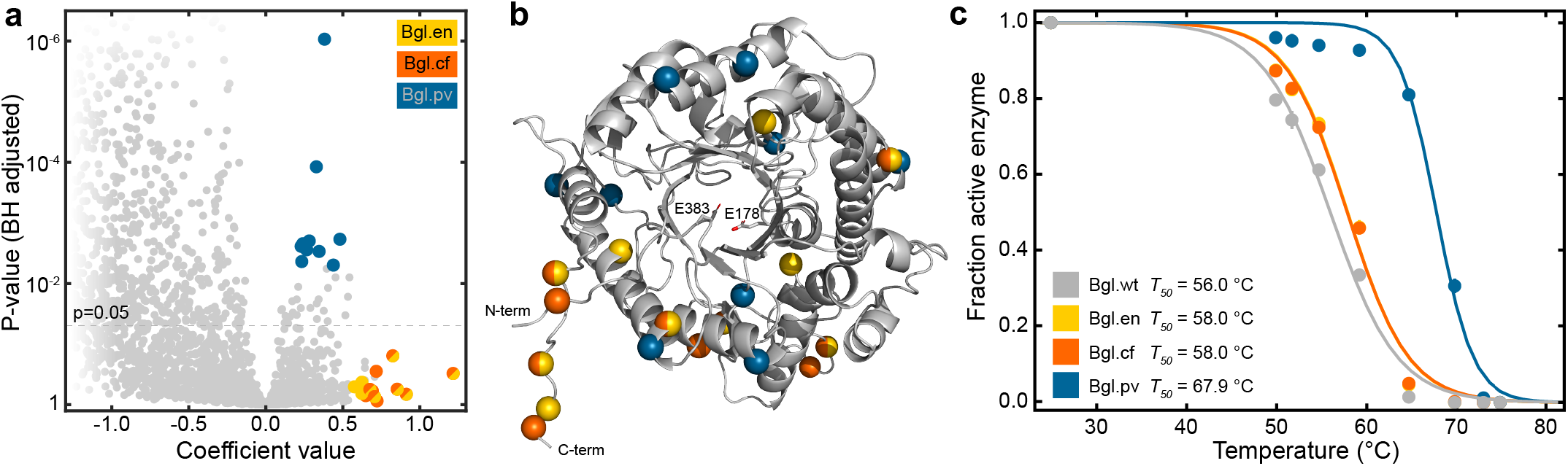
Applying the PU model to design enhanced proteins. (a) A plot of model coefficients versus p-values. Sequences were designed to combine ten mutations with the largest coefficient values, smallest p-values, or largest enrichment scores. (b) The positions chosen by the three design methods are mapped onto the Bgl3 protein structure. The structure is based on the Bgl3 crystal structure (PDB ID: 1GNX) and missing termini/loops were built in using MODELLER (Sali & Blundell 1993). (c) Thermostability curves for wild-type Bgl3 and the three designed proteins. T50 values were estimated by fitting a sigmoid function to the fraction of active enzyme. Note the curve for Bgl.en is shown in yellow and falls directly behind the orange Bgl.cf curve.

We experimentally characterized the thermostability of wild-type Bgl3 and the three designed enzymes. All three designed sequences were stabilized relative to wild-type Bgl3 (Figure 5c). The coefficient and enrichment-based designs displayed modest stability increases (~ 2 °C), while the p-value-based design was almost 12 °C more stable than wild-type Bgl3.

## Discussion

We have presented a novel supervised learning framework to infer the mapping from protein sequence to function from large-scale sequence-function data. We applied a positive-unlabeled (PU) learning approach to address the lack of negative sequence examples typically encountered in deep mutational scanning data. Our PU learning method models a protein’s true functional response as an unobserved latent variable, and then estimates how sequence maps to this latent response by maximizing the observed likelihood. Importantly, our approach leverages established statistical methods and hypothesis testing to evaluate the significance of sequence features and predictions. The PU models displayed excellent predictive ability and robustness across ten diverse protein data sets. The learned model parameters capture important aspects of protein structure and function, and can be used to design new and enhanced proteins.

We compared the PU model’s predictive ability to established structure-based and unsupervised learning methods including Rosetta, EVmutation, and DeepSequence. This is a rather unequal comparison because the PU model is trained directly on the DMS data, while the other methods are trained on peripherally related sequence/structure data. As expected, the PU model displayed substantially higher predictive performance than structure-based or unsupervised methods. These other methods have the distinct advantage that they can make reasonable predictions in the absence of DMS data, while our PU model requires DMS data for training. The relative performance of these various predictive methods is likely dependent on the particular protein activity that’s being modeling. We expect Rosetta to capture protein activities related to folding and stability; while EVmutation and DeepSequence may capture preservation of native function and the associated biophysical properties. Along these lines, supervised methods that learn the mapping to a particular property are required to model and predict non-natural protein properties. The ability to predict non-natural properties is essential for designing new proteins with behaviours beyond naturally evolved biological function.

There is a close connection between our PU model’s parameters and the enrichment scores commonly used to evaluate DMS experiments. Both methods estimate how amino acid substitutions affect a protein’s functional response. However, there are two key differences in these estimates: (1) enrichment-based methods estimate the positive-unlabeled response, while our method directly estimates the positive-negative response, and (2) enrichment-based methods estimate marginal amino acid effects and therefore make strong assumptions about the independence between sequence positions. In theory, the parameters estimated using our PU learning method should provide a more accurate and less biased estimate of how amino acid substitutions affect function. We found the PU model had greater predictive ability than enrichment on all ten protein data sets tested. While the differences in predictive performance were small (AUC differences < 0.02), these differences were statistically significant (*p*< 10^−9^) in all cases. These results suggest the learned PU model is better overall, but the predictions may not be much different from enrichment. The greatest advantage of the PU model over enrichment is the ability to perform statistical hypothesis testing to evaluate the significance of the model parameters. Hypothesis testing allows us to have confidence in the parameter estimates and predictions, and is thus essential for protein design.

We applied the learned PU models to design beta-glucosidases with improved thermal tolerance. We compared design strategies based on enrichment, PU model coefficients, and p-values. We found that enrichment and coefficient-based methods chose similar substitutions, and the resulting designs had modest increases in thermostability. In contrast, the p-value-based design contained a distinct set of substitutions and was significantly stabilized relative to the wild-type parent sequence. These results suggest that it’s better to design sequences containing high confidence substitutions rather than including uncertain substitutions with the largest magnitudes. In principle, this protein engineering strategy could be implemented iteratively, where a DMS data set is generated from an initial parent sequence, these data are used to design an improved sequence, then a new DMS data set is generated around this improved sequence, and the process is repeated. This iterative sequence optimization approach is similar to directed evolution, however, it fully leverages sequence-function information at each generation, allowing it to take larger jumps in protein sequence space.

There are several interesting extensions of the PU learning framework presented here. In this work we only considered a linear sequence-function mapping. In theory, our modeling framework could be extended to include pairwise or even higher order interactions between residues. These models would account for epistatic interactions between sites, and could possibly be used to determine contacting residues in the protein’s three-dimensional structure. We performed preliminary tests to evaluate whether we could model interactions in DMS data, and found the massive increase in system variables made the computations intractable in most cases. Future work could explore more efficient algorithms for learning from high-dimensional interaction models. Another interesting area to explore is multi-response models that consider several protein properties simultaneously. For example, we could model the Bgl3 room temperature and high temperature data sets simultaneously to directly resolve the residues responsible for protein stability. Finally, our PU modeling approach used a point estimate for a hyperparameter *π*. A more integrated modeling framework could account for uncertainty in *π* estimates and how that propagates to model coefficients and p-values.

We applied our PU learning framework to model protein sequence-function relationships. In principle, similar approaches could be used to model genotype-phenotype mappings across any level of biological organization. Positive-unlabeled data arise whenever a population of genetic variants (generated via mutagenesis, crossbreeding, etc.) is passed through a phenotypic screen/selection, and the genotypes from the before/after populations are determined using high-throughput DNA sequencing. This general format has been used to experimentally map genotype-phenotype relationships for promoters/regulatory sequences (Kosuri et al. 2013, Holmqvist et al. 2013), metabolic pathways (Ghosh & Landick 2016), microbial and mammalian genomes (Ehrenreich et al. 2010, Robins et al. 2013, Price et al. 2018, Findlay et al. 2018), and microbial communities (Kehe et al. 2019, Hsu et al. 2019).

A quantitative understanding of the mapping between protein sequence and function is important for describing natural evolution, diagnosing and treating human disease, and designing customized proteins. Advances in experimental technology have enabled researchers to map sequence-function relationships on an unprecedented scale and resolution. The resulting data are challenging to analyze because they’re typically massive, high-dimensional, contain missing sequence information, and lack negative sequence examples. Our PU learning framework provides a principled way of analyzing large-scale sequence-function data to yield biochemical insights and make quantitative predictions.

## Materials and Methods

### Linear sequence-function model

We model the sequence-function mapping *f* as a linear function of amino acid sequence. Suppose we have a protein of length *L* and a protein’s activity can be described as the sum of individual amino acid contributions:

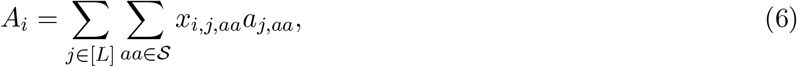

where *x*_*i,j,aa*_ ∈ {0, 1} is a binary variable that specifies whether sequence *i* has amino acid *aa* at position *j*, *a*_*j,aa*_ specifies the contribution of amino acid *aa* at position *j* to protein activity, [*L*] is the set of all positions in the sequence (1,…, *L*), and 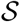 is the set of all 20 amino acids and the stop codon (*A, V*,…, *). We note that each sequence position takes on one and only one amino acid option (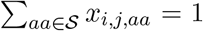 for all *i, j*) and therefore the model is over-parameterized. We introduce a reduced model:

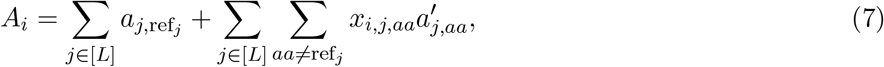

where ref_*j*_ is a reference amino acid state for the *j*th position. This reference sequence is typically the wild-type parent sequence that was used to make the DMS library. It can be shown with simple algebra that 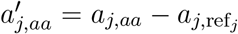, and therefore 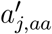 parameterizes the effect of making an amino acid substitution from the reference state ref_*j*_ to *aa* at position *j*.

Our linear sequence-function model can be specified in vector notation as:

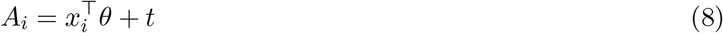

where *x*_*i*_ is a one-hot encoded vector that specifies a protein’s amino acid sequence:

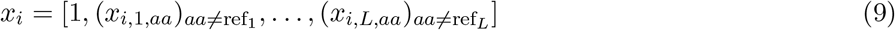

and the vector *θ* contains the model parameters:

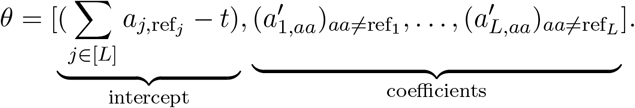

Here we see that the intercept term parameterizes the activity of the reference sequence relative to the activity threshold *t*.

### Data preprocessing

We obtained the ten large-scale sequence-function data sets from previously published work (Olson et al. 2014, Wrenbeck et al. 2019, Weile et al. 2017, Doud & Bloom 2016, Romero et al. 2015) and the Sequence Read Archive (SRA) (Leinonen et al. 2011). The details of each data set and their SRA accession numbers are available in Supplementary Table 1, and an overview of the data processing is summarized in Supplementary Figure 2. For each data set, we obtained raw FASTQ files for both unlabeled and positive sequences, and mapped these reads to a reference sequence using Bowtie2 (Langmead & Salzberg 2012). We translated the aligned gene sequences to amino acid sequences, and filtered the data sets to remove any amino acid substitutions that were observed less than ten times.

We used mode imputation to fill in any missing sequence information. Many of the analyzed data sets consisted of partial sequencing fragments (either tiled or random) because the entire gene was too long to cover with a paired-end Illumina read. The remainder of the sequence positions were unobserved. We used mode imputation to replace this missing sequence information. For nearly all DMS data sets, mode imputation simply replaces unobserved positions with the wild-type amino acid.

We converted protein sequence observations to a design matrix **X** using one-hot encoding. Each row of the design matrix **X** has the form in (9), where the reference amino acid sequence is taken to be the most frequent amino acid at each position (usually corresponding to the wild-type sequence). The DXS data set consisted of eight recombined gene fragments from one of the four DXS parent sequences (*E. coli, B. subtilis, Z. mobilis, P. trichocarpa*). These chimeric DXS sequences can be represented as an ordered sequence of “blocks” that indicates which parent the gene fragment was inherited from. We chose the *E. coli* DXS as the reference and generated dummy variables for each block change from the reference. Each data set resulted in two design matrices corresponding to unlabeled and positive sequences.

### PU model training

We trained PU models on the unlabeled and positive sequence sets for each protein data set. We computed the observed likelihood (Eqn 3) for a given data set 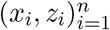 and a hyperparameter *π*. We found a stationary point of the negative observed log-likelihood using a Majorization-Minimization (MM) algorithm (Ortega & Rheinboldt 1970, Lange et al. 2000). Specifically, starting from an initial parameter value which corresponds to the null model (no features in the model), we obtained a quadratic majorizer of the negative log-likelihood function at the current parameter value and updated the current parameter with a minimizer of the quadratic majorizer function. Since the majorizer function is greater than the negative likelihood at all points, the minimizer decreases the function value of the negative likelihood compared to the function value evaluated at the current parameter value, i.e. the minimizer increases the likelihood value. This process was repeated until convergence. For the implementation of this process, we have used the **PUlasso** R package from the Comprehensive R Archive Network (CRAN) (Song & Raskutti 2018), setting the regularization parameter λ = 0 to fit the un-penalized model.

The hyperparameter *π* was either determined experimentally or tuned to maximize the model’s classification performance. For hyperparameter tuning, we used twenty log-spaced *π* values ranging from 10^−3^ to 0.5. For each *π* value, we trained a model on 90% of the data set, used the model to make predictions on the remaining 10%, and generated a receiver operating characteristic (ROC) curve for the predictions using the labeled/unlabeled response *Z*. Importantly, *π* sets an upper limit on this labeled/unlabeled ROC curve (Supplementary Figure 1), and in some instances the observed ROC curve exceeded this upper limit. These values of *π* were determined to be infeasible because they resulted in true positive rates greater than the oracle classifier. We selected the *π* value which resulted in the highest ROC-AUC value among feasible *π* values.

### Aggregating models from multiple replicates

In some instances we had data from multiple replicates that needed to be combined. For example, the Bgl3 high-temp data set had two experimental replicates (*R* = 2). We trained models on each individual replicate and then aggregated these results into a single model with estimated coefficients 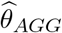 and variance-covariance matrix 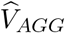. Let 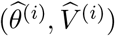 be the estimated coefficients and the variance-covariance matrix of the coefficients from the ith replicate. Here 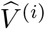 was computed as an inverse of the estimated Fisher information at 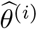. The aggregated coefficient and variance-covariance matrix were calculated as follows:

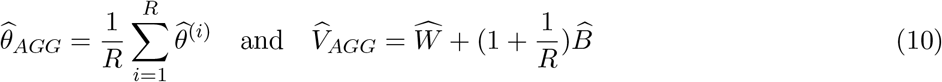

where

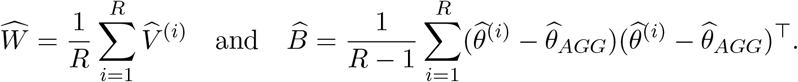

We note that the aggregated variance matrix 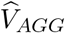 has two components: 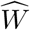 for the variation in 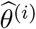 within each replicate and 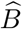 for the variation across different replicates, i.e. 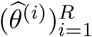. Thus the form of 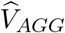 in Eqn 10 is a classical variance decomposition with the extra 1/*R* factor to account for the finite *R* (see e.g. Carpenter & Kenward (2013)).

### Evaluating and comparing model predictive ability

We used the area under the receiver operating characteristic curve (ROC-AUC) to evaluate the predictive ability of each model. With PU data, we don’t have negative examples and therefore we can’t directly calculate a model’s false positive rate (FPR). Instead, we used the labeled-unlabeled response (*Z*) to calculate the false positive rate (FPR^*PU*^) and true positive rate (TPR^*PU*^), and then performed a correction (Jain et al. 2017) to obtain ROC curves and ROC-AUC values:

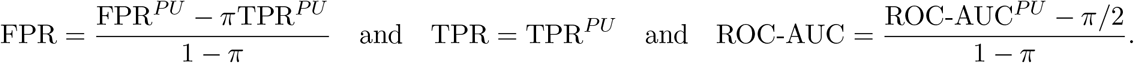

This PU ROC curve correction is illustrated in Supplementary Figure 1. We obtained corrected ROC curves and ROC-AUC values for each of the ten cross-validation folds, and then averaged over folds to obtain the model’s corrected ROC curve and ROC-AUC value.

We used a corrected repeated cross-validation test to compare the predictive ability between the PU model and site-wise enrichment (Bouckaert & Frank 2004). Importantly, this test controls inflated type 1 error caused by data overlaps in cross-validation folds (Dietterich 1998, Nadeau & Bengio 2003) and also has high replicability (BoucaWe used 10-fold cross-validation to evaluatekert & Frank 2004). The test involves running *K*-fold cross validation for *R* independent runs and comparing models using a corrected test statistic. For each run *i* = 1,…, *R*, we split the data randomly into *K* sub-samples and fit one model for each fold *j* = 1,…, *K*. We used *R* = *K* = 10 as recommended by the authors (Bouckaert & Frank 2004). Let ROC-AUC(M)_*ij*_ be the corrected ROC-AUC value for model *M* from the *i*th run and *j*th cross-validation fold. For the enrichment-based predictions, we used an additive model that summed all individual enrichment scores. We define the difference between the models *d*_*ij*_ ≔ ROC-AUC(PU model)_*ij*_ − ROC-AUC(enrich)_*ij*_ and the standard deviation of this difference 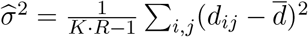. The test statistic *t* was calculated as follows

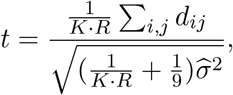

and was compared with the *t* distribution with *K* · *R* − 1 degrees of freedom.

### Predictions using Rosetta, EVmutation, and DeepSequence

We made protein function predictions using established structure-based and unsupervised learning methods including Rosetta, EVmutation, and DeepSequence (Alford et al. 2017, Hopf et al. 2017, Riesselman et al. 2018). For Rosetta modeling, we searched the Protein Data Bank to identify the structure most similar to the DMS data’s reference sequence and used this as a template for Rosetta comparative modeling using the default options (Song et al. 2013). We sampled 500 random sequences from both the unlabeled and positive sequence sets, built Rosetta models for these 1000 sequences, and calculated the Rosetta energy for each. We used these calculated Rosetta energies to create ROC curves classifying unlabeled and positive sets, and corrected these ROC curves to account for PU data.

We made predictions using EVmutation and DeepSequence models for each protein data set. We created family multiple sequence alignments (MSAs) using jackhmmer (Wheeler & Eddy 2013) to query the DMS data set’s reference sequence against the UniRef90 sequence database (Suzek et al. 2015). For DXS, we chose the *E. Coli* parent as the jackhmmer reference sequence. For Rocker, we had to relax the inclusion threshold (jackhmmer domE option set 10000) to include additional sequences because Rocker is a *de novo* designed protein. We filtered the jackhmmer results to remove amino acid insertions relative to the reference sequence and also removed any resulting sequences that had less than 50% coverage over the reference sequence. We trained EVmutation and DeepSequence models on these curated MSAs using the default options. For EVmutation, we scored all sequences in each data set, except for GB1, where we sampled 10^6^ random sequences and DXS, where we sampled 10^4^ random sequences. For DeepSequence, we scored 10^4^ random sequences from each data set. We used the EVmutation and DeepSequence scores to create corrected ROC curves for each data set.

### Statistical hypothesis testing

We performed hypothesis tests to determine which features “significantly” affect protein function. We calculated the *Z* statistic *z*_*j*_ to test whether a feature *j* affects protein function or not (i.e. *H*_0_ : *θ*_*j*_ = 0):

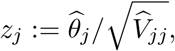

where 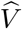 is the estimated variance-covariance matrix of 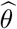, computed as the inverse of the estimated Fisher information. We obtained p-values under the null hypothesis that *θ*_*j*_ = 0 and computing tail probabilities:

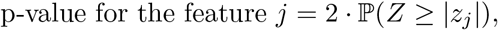

where *Z* is a standard normal variable. These p-values were then adjusted using the Benjamini-Hochberg (BH) procedure to account for multiple hypothesis testing and control the false discovery rate (Benjamini & Hochberg 1995). We considered a feature to be significant if its BH-adjusted p-value was greater than 0.05.

### Evaluating PU model stability

We used 10-fold cross-validation to evaluate the stability of the fitted PU model’s parameter estimates and selected features. We calculated the coefficient of variation for each feature *j* across different cross-validation folds:

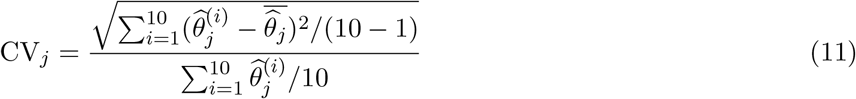

where 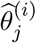 is an estimated coefficient for the *j*th feature from the *i*th cross-validation fold. The absolute value of this coefficient of variation is a measure of a coefficient’s relative variability (Supplementary Figure 3d).

We evaluated feature selection stability by comparing the set of selected features across different cross-validation folds. We defined a selection stability measure (SS) as the average fraction of common selected features:

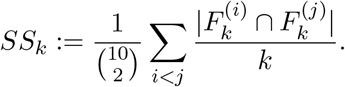

where *k* is the selection size, *i* and *j* are different cross-validation folds, and 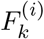 is the set containing features with the *k* smallest p-values from the ith fold. We computed *SS*_*k*_ for *k* = 1,…, *K*, where *K* was chosen to be the number of the significant features (BH-adjusted *p* < 0.05). We then averaged *SS*_*k*_ over *k* to obtain the average feature selection stability 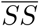. A value of 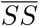 that approaches one indicates that many common features are selected across different cross-validation folds (Supplementary Figure 3d).

### Beta-glucosidase cloning, expression, and characterization

We designed the genes encoding the Bgl3 variants by making codon substitutions into a base Bgl3 gene sequence. If there were multiple codon options for an amino acid, we chose the particular codon randomly from a set of 31 codons that are optimized for expression in E coli (Böel et al. 2016). We ordered the designed genes as gBlocks from IDT and cloned them into a protein expression vector (pET22b) using GoldenGate cloning. We verified the sequences of all genes using Sanger sequencing with the T7 promoter and T7 terminator primers.

We expressed the Bgl3 variants shaking at 30 °C in a 5-mL MagicMedia (Invitrogen) culture overnight. We then pelleted the expression culture by centrifugation and froze at −20 °C. We resuspended the cell pellets in lysis buffer [0.3× BugBuster (Novagen), 30 kU/mL rLysozyme (Novagen), and 50 U/mL DNase I (New England Biolabs) in 100 mM potassium phosphate, pH 7.2] and performed serial dilutions to determine the linear range of the enzyme assay. We then diluted all samples in 100 mM potassium phosphate, pH 7.2 to be within the linear range and have similar end-point activities.

We arrayed the diluted cell extracts into 96-well PCR plates and heated the samples over multiple temperatures (50–75 °C) for 15 min using a gradient thermocycler. After the heat step, we quantified the remaining functional enzyme by adding the fluorogenic substrate 4-methylumbelliferyl-*β*-D-glucopyranoside (Sigma) to a final concentration of 1 mM. We monitored the reaction progress by fluorescence spectroscopy (372 nm excitation/445 nm emission), and determined the rate by fitting a linear function to the progress curves. We normalized all rates to enzyme samples that had been incubated at room temperature (25 °C). The *T*_50_ (temperature where 50% of the protein is inactivated in 15 min) was determined by fitting a shifted sigmoid function to the thermal inactivation curves. All measurements were performed in at least duplicate with the mean *T*_50_ values reported.

## Acknowledgments

We would like to acknowledge funding support from the NIH grants R35 GM119854 and R01 GM131381, and the NSF grant DMS-1811767.

## Author Contributions

H.S., G.R., and P.A.R. conceived the project. H.S. and G.R. developed the PU learning methods and code. H.S., E.C.H., and P.A.R analyzed the data. B.B. performed the Rosetta, EVmutation, and DeepSequence analysis. B.B. performed all experimental work and analysis. H.S. and P.A.R. wrote the paper with feedback from all other authors.

## STAR Methods

### Enrichment-based methods provide attenuated estimates of a mutation’s effect due to latent positive sequences

Here we demonstrate that enrichment scores calculated from positive-unlabeled (PU) responses provide *biased* estimates of a mutation’s effect. Consider the population enrichment score 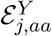 calculated from the positive-negative (PN) responses:

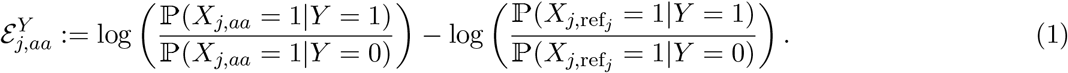

which correspond to an amino acid substitution’s true marginal effect, i.e. the effect of changing site *j* from ref_*j*_ to amino acid *aa* while allowing all other positions to vary. Also consider the population enrichment score 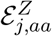 calculated from PU responses

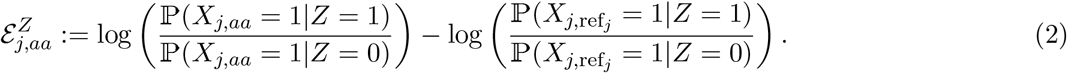

We consider 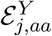 to be the true marginal effect, and show that enrichment calculated from the PU response 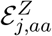 is not equivalent (i.e. 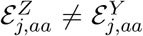). We also demonstrate that 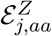 provides an attenuated estimate of the true mutational effect due to latent positive sequences in the unlabeled set.

Since labeled sequences are positive, for any *k* ∈ (*A, V*,…, *) we have

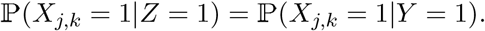

On the other hand,

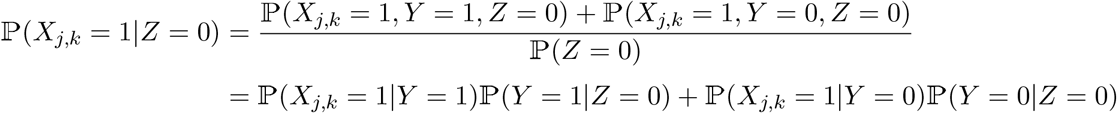

Therefore,

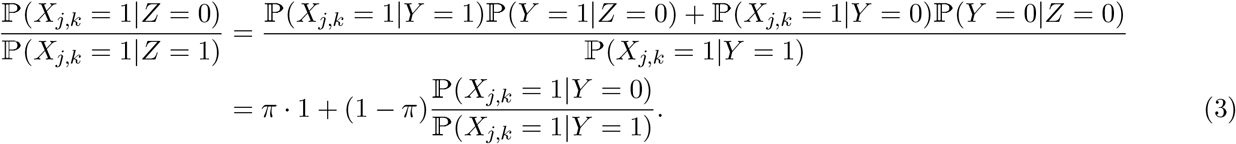

where 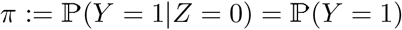. Applying (3) to (2) with *k* ∈ {ref_*j*_, *aa*},

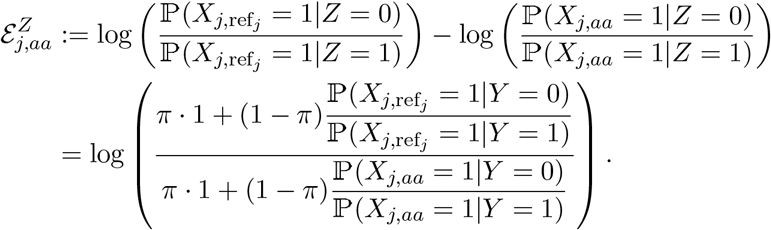

To ease notation, let

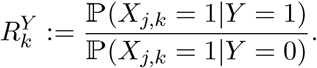

Then,

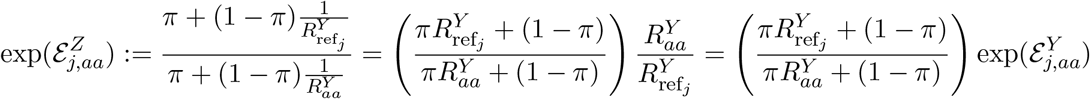

Note 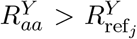 if and only if 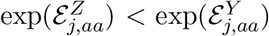. In other words, 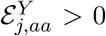 if and only if 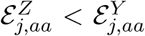. Since 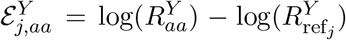 by definition, 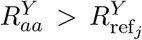 is equivalent to 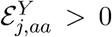. That is, 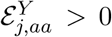 if and only if 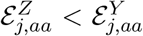. Therefore, 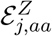 underestimates the effect amino acid substitutions with positive effects 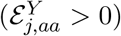 and overestimates substitutions with negative effects.

### Site-wise enrichment captures marginal effects, which are biased

Here we demonstrate that site-wise enrichment scores capture marginal mutational effects that are in general biased for estimating an amino acid substitution’s true effect 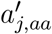. Suppose the true positive-negative responses (*y*_*i*_) were available. Enrichment scores can be calculated from this positive-negative (PN) response as:

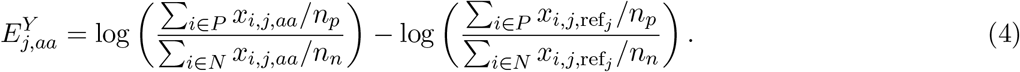

where *P* and *N* are the positive and negative sets of sequences. We demonstrate that 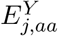 are in general biased for 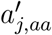.

Recall the logistic model from Eqn 2 in the main text:

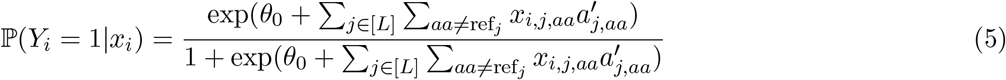

where 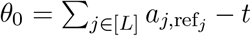 and 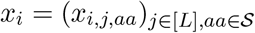 is a one-hot encoded vector of the sequence *i*. Maximizing the likelihood function 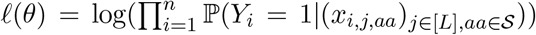 will produce consistent estimates for 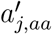. Consistent estimates will approach the true value of 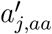 as the number of data points increases. It is a well-known result in Statistics that the maximum likelihood estimator provides a consistent estimate.

Site-wise enrichment scores calculated from the true PN response (*y*_*i*_) are equivalent to the maximum likelihood estimates (MLE) from a logistic model that considers only one site at a time:

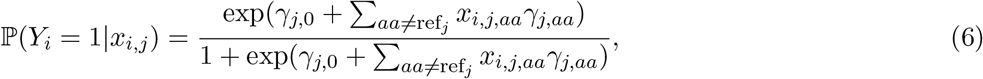

where 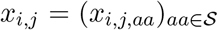. Note how this equation only includes terms related to the *j*th position, in contrast to Eqn 5, which sums over all *j* ∈ [*L*].

We demonstrate that the MLE of 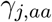 is equal to the enrichment score calculated from the true positive-negative responses 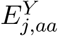. The maximum likelihood estimate 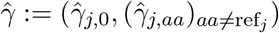 solves likelihood equations

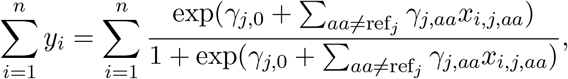

and for all *aa* ≠ ref_*j*_,

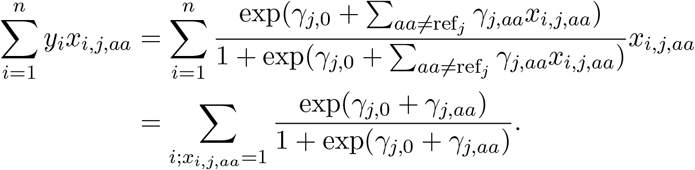

That is,

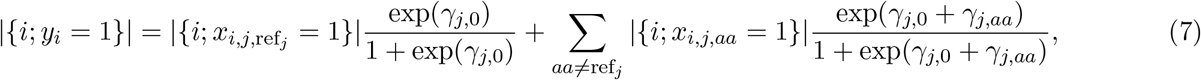

and for all *aa* ≠ ref_*j*_,

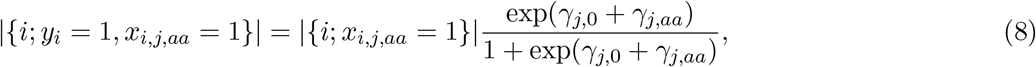

where |*A*| denotes the size of *A*. Solving Eqn 7 and 8 for 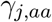,

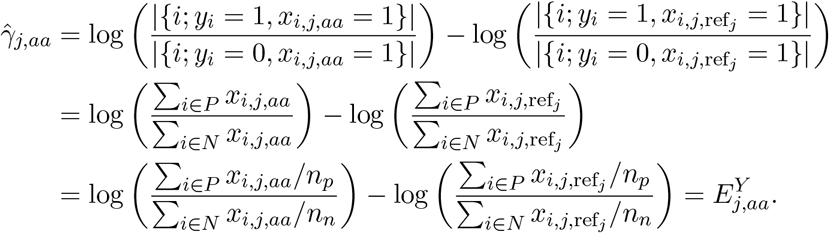

Thus 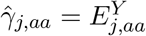.

In addition, 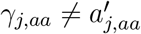 in general because the estimate for 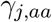 does not consider other sites and is therefore subject to omitted variable bias (e.g. see [1]). From this we can conclude that site-wise enrichment scores calculated from the true positive-negative response 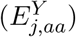 provide biased estimates for 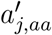.

There exists one exception when sequence positions are independent given each response class. In this case, 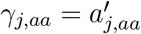 and enrichment scores from the positive-unlabeled response provide consistent estimates 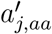. Class conditional independence between positions is unlikely to hold for any DMS data set due to biophysical interactions between sites, correlated sequence variables in multi-mutant sequences, and the nonlinear threshold in the response.

## Supplementary Figures

**Supplementary Figure 1:**
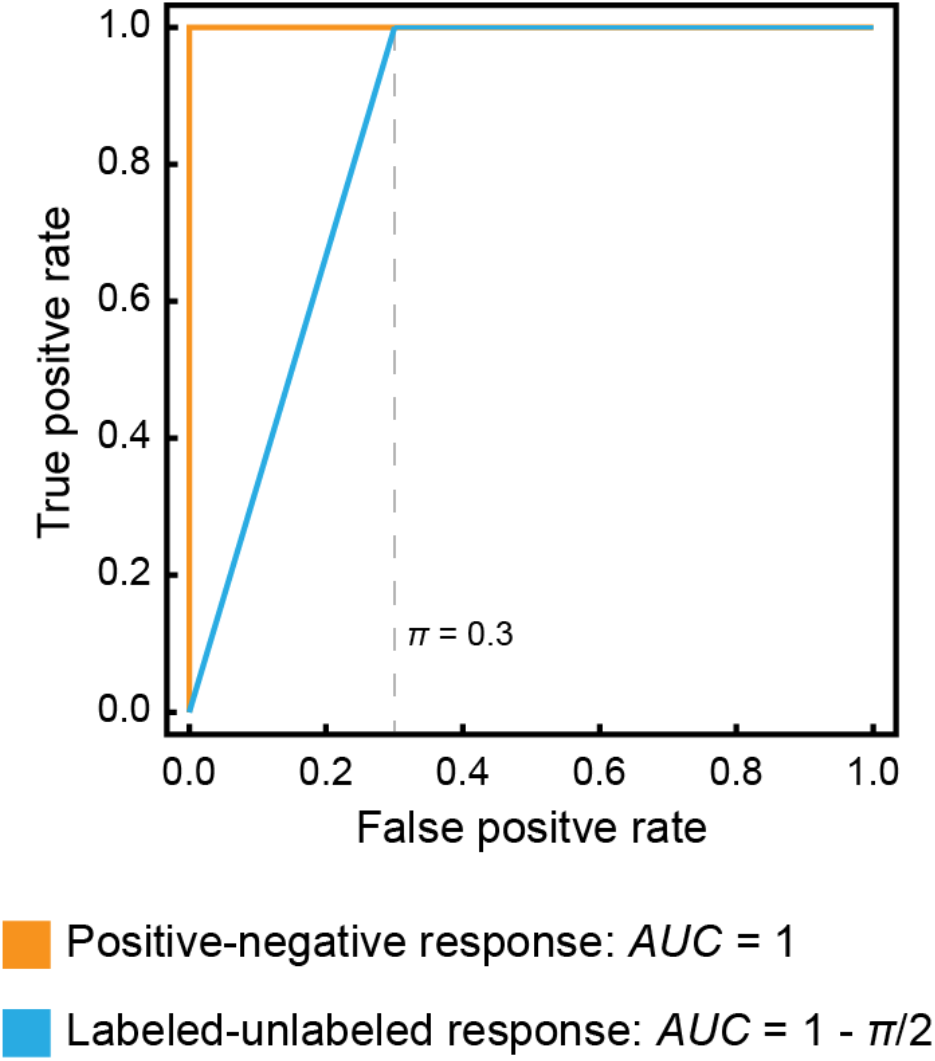
Illustration of the ROC curves for the oracle (perfect) classifier using the positive-negative response (*Y*) or the labeled-unlabeled response (*Z*). Note the labeled-unlabeled response is identical to the positive-unlabeled response because all labeled sequences are positive. The oracle classifier cannot perfectly separate the labeled (positive) and unlabeled classes because the unlabeled class contains positive examples. The value of *π* (0.3 in this example) sets an upper limit on the performance of any labeled-unlabeled classifier. This upper limit can be used to correct ROC curves generated from positive-unlabeled data to estimate the true positive-negative-ROC curve.

**Supplementary Figure 2:**
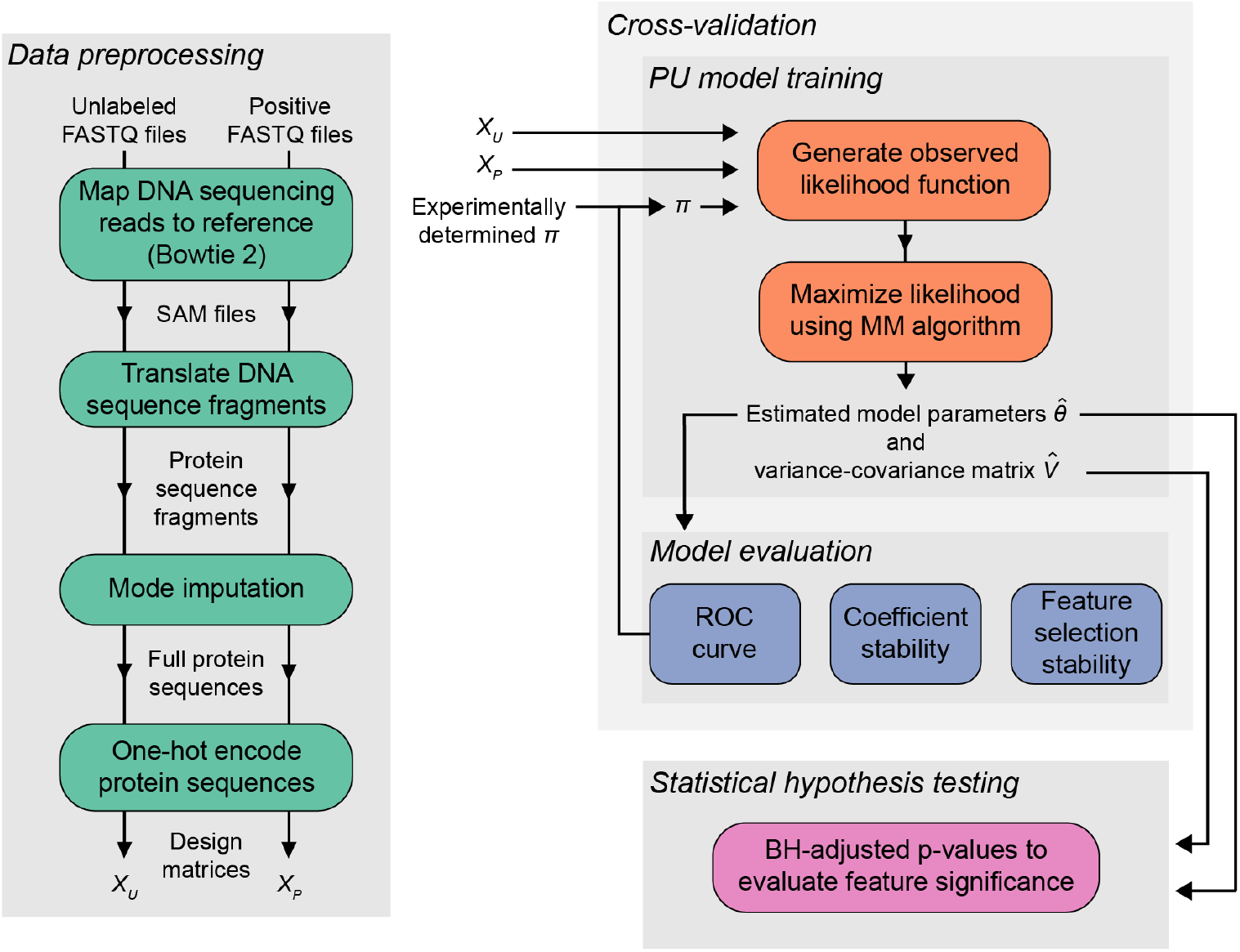
An overview of the data processing and analysis workflow.

**Supplementary Figure 3:**
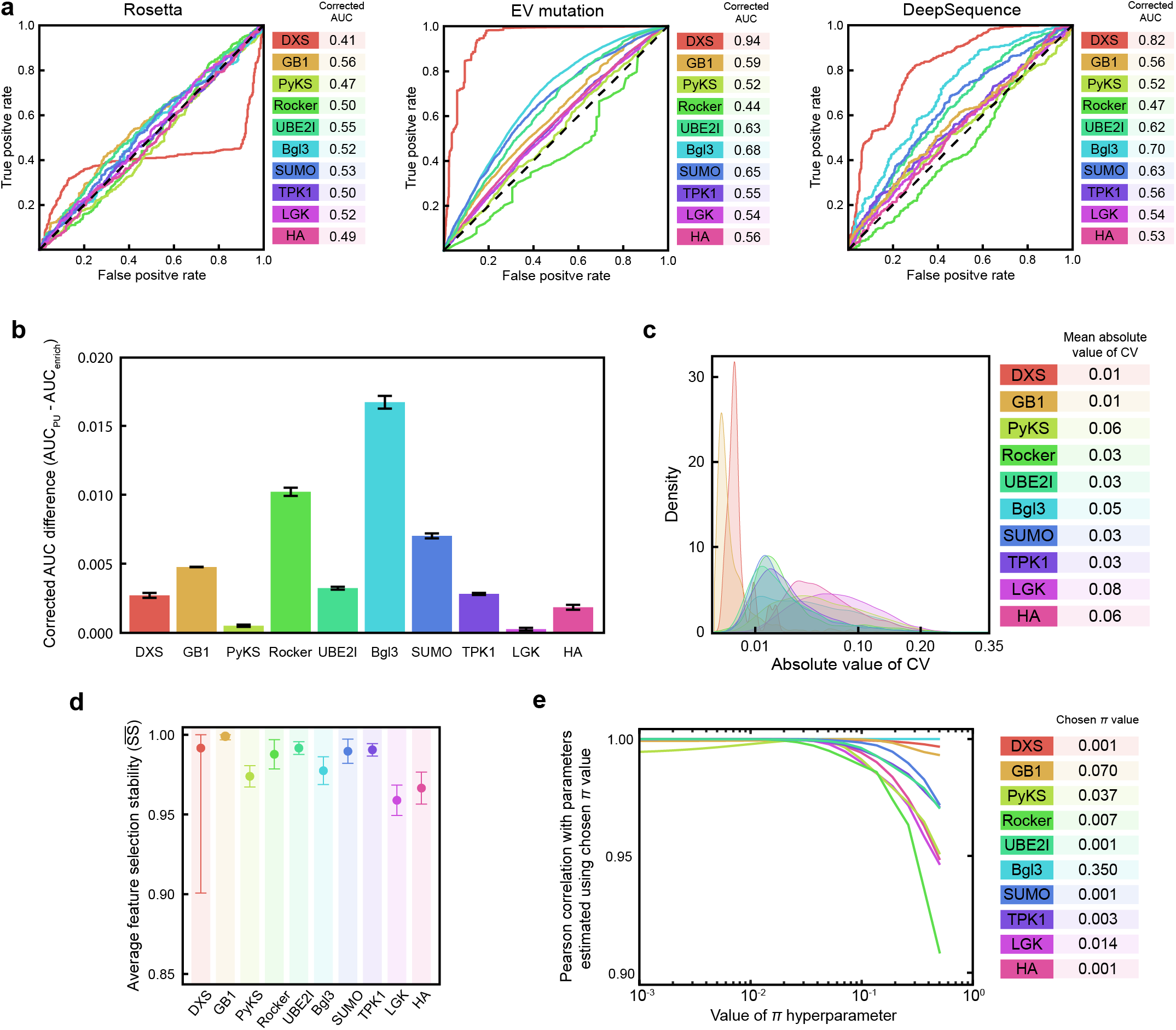
(a) Receiver operating characteristic (ROC) curves for structure-based and unsupervised learning methods. ROC curves were corrected to account for PU data using the same *π* as the PU model to allow direct comparison. (b) The PU model displayed higher AUC values than site-wise enrichment for all ten data sets. These AUC differences were small, but statistically significant, with *p* < 10^−9^ for all data sets. (c) Density curves of the model coefficient’s absolute coefficient of variation (CV). The mean absolute CVs range from 0.01 to 0.08, indicating the model coefficients are highly robust to cross-validation. (d) The average value of the feature selection stability 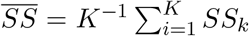, where *SS*_*k*_ is measured as the average fraction of common features with selection size *k* across 10 cross-validation folds and *K* is the number of significant features. On average, over 95% of the selected features overlap, indicating the model’s feature ranking was stable across different cross-validation folds. (e) PU models are highly insensitive to the particular choice of the *π* hyperparameter. We trained models using forty log-spaced *π* values ranging from 10^−3^ to 0.5, and calculated the Pearson correlation between the chosen model’s coefficients and all other models. The *π* value for the chosen model is indicated in the table. We see models generated across the entire range of *π* show very strong correlation with the chosen model. This indicates that the model interpretation, conclusions, and predictions are insensitive to our estimate for *π*.

**Supplementary Figure 4:**
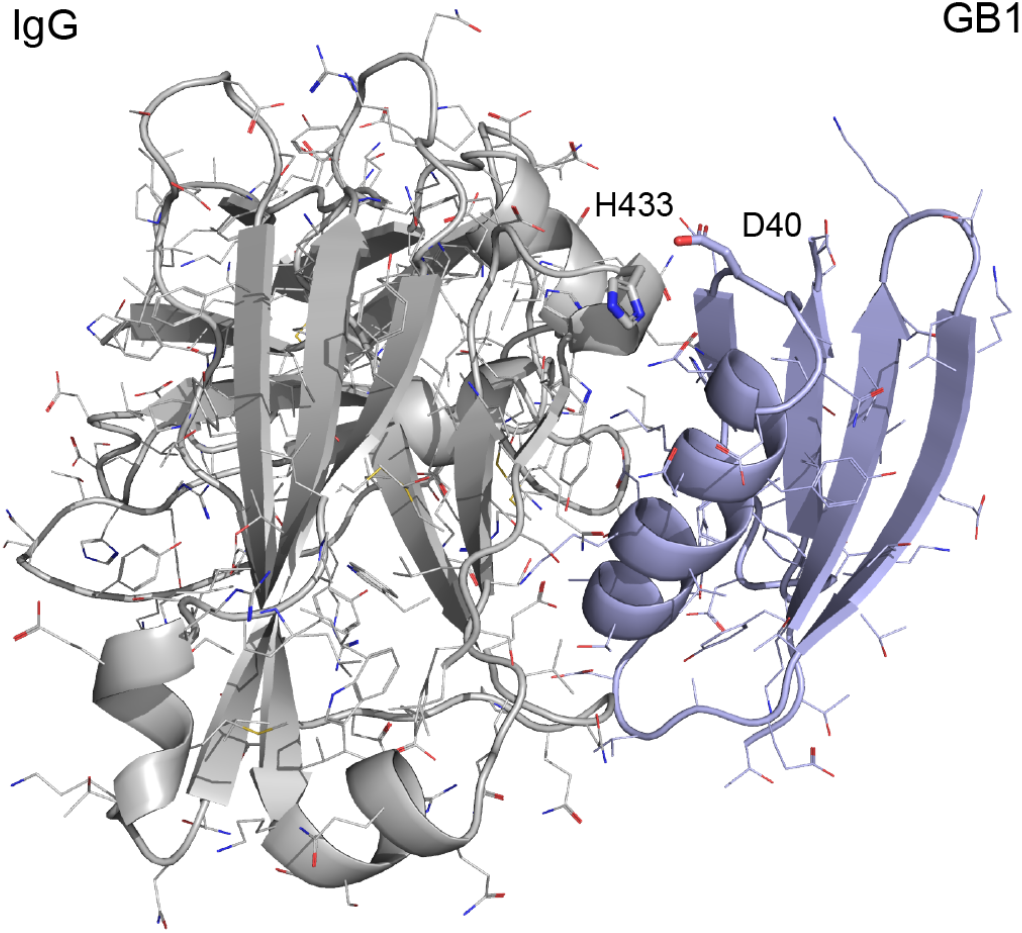
Substituting D40 for aromatic residues could result in potential pi-pi or cation-pi interactions with IgG’s H433. In the crystal structure (PDB ID: 1FCC), IgG is shown in grey and GB1 in light blue.

**Supplementary Figure 5:**
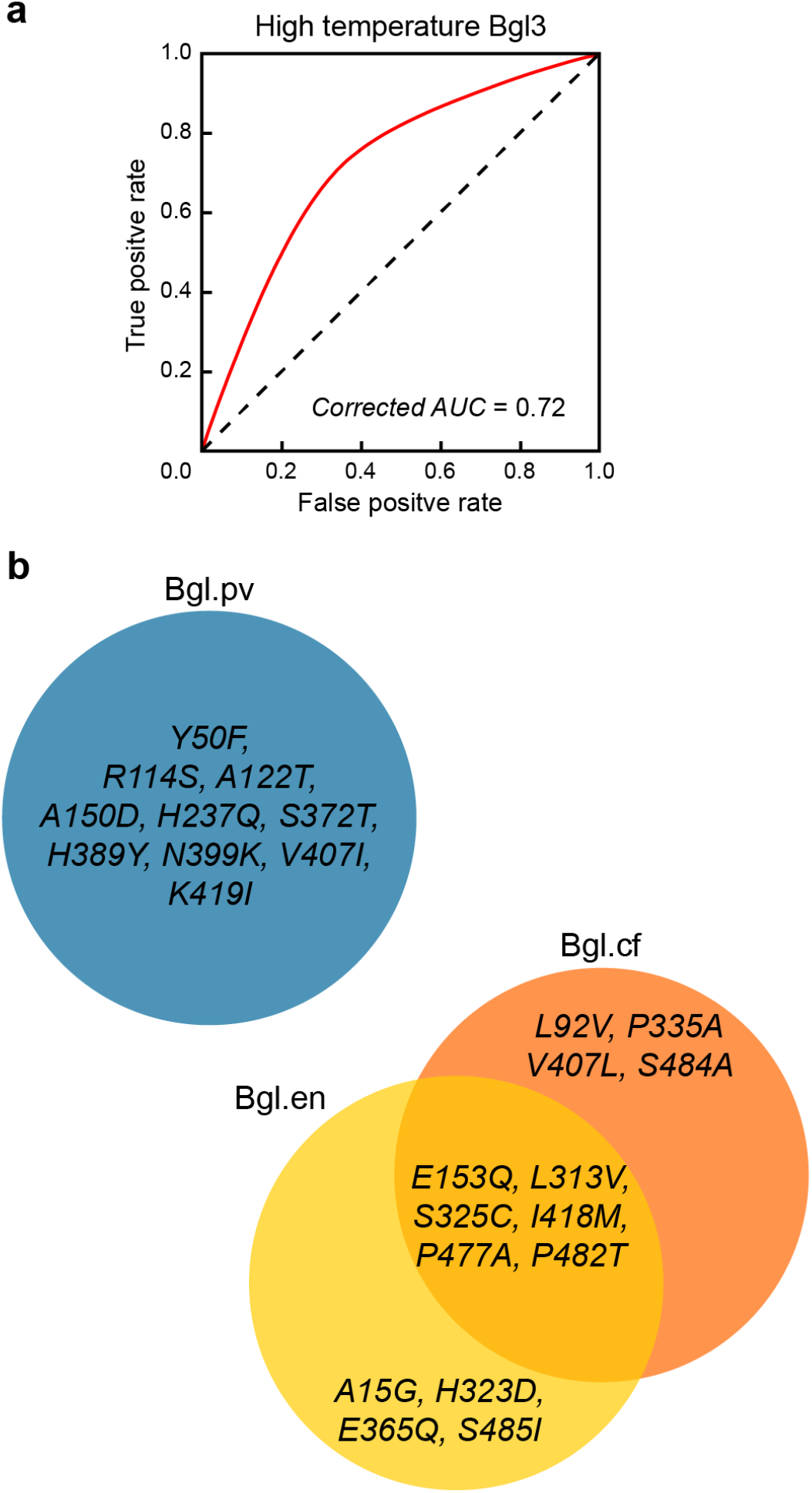
(a) Corrected ROC curve generated for the high-temperature Bgl3 data set. (b) Venn diagram depicting the amino acid substitutions present in the three designed beta-glucosidases. Each design has ten substitutions from wild-type Bgl3. The enrichment- and coefficient-based designs (Bgl.en and Bgl.cf, respectively) share six common substitutions.

| **Data set name** | **Reference PubMed ID** | **Protein name** | **Reference Sequence UniProt ID** | **Sequence length** | **Library type** |
| --- | --- | --- | --- | --- | --- |
| GB1 | 25455030 | Protein G domain B1 | P06654 | 55 | Double site saturation mutagenesis |
| PyKS | 30721031 | Pyrrolidine Ketide Synthase | A0A3G4RHW | 393 | Single site saturation mutagenesis |
| LGK | 30721031 | Levoglucosan kinase | B3VI55 | 445 | Single site saturation mutagenesis |
| SUMO | 29269382 | Small ubiquitin-related modifier 1 | P63165 | 98 | Error-prone PCR |
| UBE2I | 29269382 | SUMO-conjugating enzyme UBC9 | P63279 | 167 | Error-prone PCR |
| TPK1 | 29269382 | Thiamin pyrophosphokinase 1 | Q9H3S4 | 243 | Error-prone PCR |
| HA | 27271655 | Hemagglutinin | P03454 | 565 | Random codon mutagenesis |
| Bgl3 | 26040002 | Beta-glucosidase | Q59976 | 501 | Error-prone PCR |
| Bgl3-HT | 26040002 | Beta-glucosidase | Q59976 | 501 | Error-prone PCR |
| DXS | To be publish | 1-deoxy-D-xylulose-5-phosphate synthase | P77488 | 669 | Gene shuffling |
| Rocker | To be publish | Rocker | None | 25 | Double site saturation mutagenesis |

| Target function | Screening/selection method | Sequencing method | Number of unlabeled seqs (millions) | Number of positive seqs (millions) | Imputation method | Value of $\pi$ hyper-parameter |
| --- | --- | --- | --- | --- | --- | --- |
| Binding | mRNA display | Illumina HiSeq 2x100 bp | 323.1 | 259.5 | None | 0.070 |
| Enzyme folding | FACS screening | Illumina MiSeq 2x250 bp | 1.5 | 0.9 | Mode | 0.037 |
| Enzyme folding | FACS screening | Illumina MiSeq 2x250 bp | 0.8 | 0.7 | Mode | 0.014 |
| Post-translational modification | Growth selection | Illumina NextSeq 2x150 bp | 2.5 | 2.3 | Mode | 0.001 |
| Enzyme activity | Growth selection | Illumina NextSeq 2x150 bp | 4.1 | 3.5 | Mode | 0.001 |
| Enzyme activity | Growth selection | Illumina NextSeq 2x150 bp | 6.9 | 3.6 | Mode | 0.003 |
| Receptor binding | Viral infection | Illumina HiSeq 2x250 bp | 1.8 | 1.3 | Mode | 0.001 |
| Enzyme activity | Microfluidic screening | Illumina MiSeq 2x300 bp | 1.5 | 2.6 | Mode | 0.350 |
| Enzyme activity after heating | Microfluidic screening | Illumina MiSeq 2x300 bp | 1.9 | 3.7 | Mode | 0.200 |
| Enzyme activity | Growth selection | Oxford Nanopore GridION | 0.7 | 0.7 | None | 0.001 |
| Membrane transport | FACS screening | Illumina MiSeq 2x150 bp | 1.3 | 0.7 | None | 0.007 |

| Method for<br>determining $\pi$ | Unlabeled SRA accessions | Positive SRA accessions |
| --- | --- | --- |
| Hyperparam. search | SRR2054943 | SRR2054944 |
| Hyperparam. search | SRR8238223,SRR8238221,Sf | SRR8238222,SRR8238220,SRR8238218,SRR8238216 |
| Hyperparam. search | SRR8238225,SRR8238229,Sf | SRR8238224,SRR8238228,SRR8238230,SRR8238226 |
| Hyperparam. search | SRR5680627,SRR5680629,Sf | SRR5680625,SRR5680634,SRR5680638 |
| Hyperparam. search | SRR5680697,SRR5680703,Sf | SRR5680699,SRR5680705,SRR5680707,SRR5680727,SRR5680733 |
| Hyperparam. search | SRR5680677,SRR5680635,Sf | SRR5680669,SRR5680632,SRR5680722,SRR5680719,SRR5680689,SRR5680692,SRR5680 |
| Hyperparam. search | SRR3113656 | SRR3113660 |
| Experimental | SRR11472224 | SRR11472223, SRR11472221, SRR11472220 |
| Experimental | SRR11472219, SRR11472215 | SRR11472218, SRR11472216, SRR11472214, SRR11472217, SRR11472222 |
| Hyperparam. search | SRR11467347 | SRR11467346 |
| Hyperparam. search | Pending | Pending |

## References

Abriata, L. A., Bovigny, C. & Dal Peraro, M. (2016), ‘Detection and sequence/structure mapping of biophysical constraints to protein variation in saturated mutational libraries and protein sequence alignments with a dedicated server’, BMC Bioinformatics 17(1).

Alford, R. F., Leaver-Fay, A., Jeliazkov, J. R., O’Meara, M. J., DiMaio, F. P., Park, H., Shapovalov, M. V., Renfrew, P. D., Mulligan, V. K., Kappel, K., Labonte, J. W., Pacella, M. S., Bonneau, R., Bradley, P., Dunbrack, R. L., Das, R., Baker, D., Kuhlman, B., Kortemme, T. & Gray, J. J. (2017), ‘The Rosetta All-Atom Energy Function for Macromolecular Modeling and Design’, Journal of Chemical Theory and Computation 13(6), 3031–3048.

Alvizo, O., Nguyen, L. J., Savile, C. K., Bresson, J. A., Lakhapatri, S. L., Solis, E. O. P., Fox, R. J., Broering, J. M., Benoit, M. R., Zimmerman, S. A., Novick, S. J., Liang, J. & Lalonde, J. J. (2014), ‘Directed evolution of an ultrastable carbonic anhydrase for highly efficient carbon capture from flue gas’, Proceedings of the National Academy of Sciences 111(46), 16436–16441.

Bedbrook, C. N., Yang, K. K., Robinson, J. E., Mackey, E. D., Gradinaru, V. & Arnold, F. H. (2019), ‘Machine learning-guided channelrhodopsin engineering enables minimally invasive optogenetics’, Nature Methods 16(11), 1176–1184.

Benjamini, Y. & Hochberg, Y. (1995), ‘Controlling the false discovery rate: A practical and powerful approach to multiple testing’, J. R. Stat. Soc. Series B Stat. Methodol. 57(1), 289–300.

Bloom, J. D. (2015), ‘Software for the analysis and visualization of deep mutational scanning data’, BMC Bioinformatics 16(1).

Böel, G., Letso, R., Neely, H., Price, W. N., Wong, K. H., Su, M., Luff, J. D., Valecha, M., Everett, J. K., Acton, T. B., Xiao, R., Montelione, G. T., Aalberts, D. P. & Hunt, J. F. (2016), ‘Codon influence on protein expression in E. coli correlates with mRNA levels’, Nature 529(7586), 358–363.

Boucher, J. I., Cote, P., Flynn, J., Jiang, L., Laban, A., Mishra, P., Roscoe, B. P. & Bolon, D. N. (2014), ‘Viewing protein fitness landscapes through a next-gen lens’, Genetics.

Bouckaert, R. R. & Frank, E. (2004), Evaluating the replicability of significance tests for comparing learning algorithms, in ‘Advances in Knowledge Discovery and Data Mining’, Springer Berlin Heidelberg, pp. 3–12.

Carpenter, J. & Kenward, M. (2013), Multiple Imputation and its Application, John Wiley & Sons.

Dietterich, T. G. (1998), ‘Approximate statistical tests for comparing supervised classification learning algorithms’, Neural Comput. 10(7), 1895–1923.

Doud, M. B. & Bloom, J. D. (2016), ‘Accurate measurement of the effects of all amino-acid mutations on influenza hemagglutinin’, Viruses 8(6).

Ehrenreich, I. M., Torabi, N., Jia, Y., Kent, J., Martis, S., Shapiro, J. A., Gresham, D., Caudy, A. A. & Kruglyak, L. (2010), ‘Dissection of genetically complex traits with extremely large pools of yeast segregants’, Nature 464(7291), 1039–1042.

Elkan, C. & Noto, K. (2008), Learning classifiers from only positive and unlabeled data, in ‘Proceedings of the 14th ACM SIGKDD International Conference on Knowledge Discovery and Data Mining’, KDD’08, ACM, New York, NY, USA, pp. 213–220.

Findlay, G. M., Daza, R. M., Martin, B., Zhang, M. D., Leith, A. P., Gasperini, M., Janizek, J. D., Huang, X., Starita, L. M. & Shendure, J. (2018), ‘Accurate classification of BRCA1 variants with saturation genome editing’, Nature 562(7726), 217–222.

Fowler, D. M. & Fields, S. (2014), ‘Deep mutational scanning: a new style of protein science’, Nature Methods 11, 801–807.

Ghosh, I. N. & Landick, R. (2016), ‘OptSSeq: High-Throughput Sequencing Readout of Growth Enrichment Defines Optimal Gene Expression Elements for Homoethanologenesis’, ACS Synthetic Biology 5(12), 1519–1534.

Holmqvist, E., ReimegÅrd, J. & Wagner, E. G. H. (2013), ‘Massive functional mapping of a 5-UTR by saturation mutagenesis, phenotypic sorting and deep sequencing’, Nucleic Acids Research 41(12).

Hopf, T. A., Ingraham, J. B., Poelwijk, F. J., Schärfe, C. P. I., Springer, M., Sander, C. & Marks, D. S. (2017), ‘Mutation effects predicted from sequence co-variation’, Nature Biotechnology 35(2).

Hsu, R. H., Clark, R. L., Tan, J. W., Ahn, J. C., Gupta, S., Romero, P. A. & Venturelli, O. S. (2019), ‘Microbial Interaction Network Inference in Microfluidic Droplets’, Cell Systems 9(3), 229–242.

Jain, S., White, M. & Radivojac, P. (2017), Recovering true classifier performance in positive-unlabeled learning, in ‘Thirty-First AAAI Conference on Artificial Intelligence’.

Jha, R. K., Gaiotto, T., Bradbury, A. R. & Strauss, C. E. (2014), ‘An improved Protein G with higher affinity for human/rabbit IgG Fc domains exploiting a computationally designed polar network’, Protein Engineering, Design and Selection 27(4), 127–134.

Kehe, J., Kulesa, A., Ortiz, A., Ackerman, C. M., Thakku, S. G., Sellers, D., Kuehn, S., Gore, J., Friedman, J. & Blainey, P. C. (2019), ‘Massively parallel screening of synthetic microbial communities’, Proceedings of the National Academy of Sciences of the United States of America 116(26), 12804–12809.

Klesmith, J. R. & Hackel, B. J. (2019), ‘Improved mutant function prediction via PACT: Protein Analysis and Classifier Toolkit’, Bioinformatics 35(16), 2707–2712.

Kosuri, S., Goodman, D. B., Cambray, G., Mutalik, V. K., Gao, Y., Arkin, A. P., Endy, D. & Church, G. M. (2013), ‘Composability of regulatory sequences controlling transcription and translation in Escherichia coli’, Proceedings of the National Academy of Sciences of the United States of America 110(34), 14024–14029.

Lange, K., Hunter, D. R. & Yang, I. (2000), ‘Optimization transfer using surrogate objective functions’, J. Comput. Graph. Stat. 9(1), 1–20.

Langmead, B. & Salzberg, S. L. (2012), ‘Fast gapped-read alignment with Bowtie 2’, Nature Methods 9(4), 357–359.

Leinonen, R., Sugawara, H. & Shumway, M. (2011), ‘The Sequence Read Archive’, Nucleic Acids Research 39(Database), D19–D21.

Liao, J., Warmuth, M. K., Govindarajan, S., Ness, J. E., Wang, R. P., Gustafsson, C. & Minshull, J. (2007), ‘Engineering proteinase K using machine learning and synthetic genes’, BMC Biotechnology 7(1), 16.

Liu, B., Dai, Y., Li, X., Lee, W. S. & Yu, P. S. (2003), Building text classifiers using positive and unlabeled examples, in ‘Third IEEE International Conference on Data Mining’, pp. 179–186.

Mazurenko, S., Prokop, Z. & Damborsky, J. (2020), ‘Machine-learning-guided directed evolution for protein engineering’, ACS Catalysis 10(2), 1210–1223.

Morcos, F., Pagnani, A., Lunt, B., Bertolino, A., Marks, D. S., Sander, C., Zecchina, R., Onuchic, J. N., Hwa, T. & Weigt, M. (2011), ‘Direct-coupling analysis of residue coevolution captures native contacts across many protein families’, Proceedings of the National Academy of Sciences 108(49), E1293–E1301.

Mordelet, F. & Vert, J.-P. (2011), ‘ProDiGe: Prioritization of disease genes with multitask machine learning from positive and unlabeled examples’, BMC Bioinformatics 12, 389.

Nadeau, C. & Bengio, Y. (2003), ‘Inference for the generalization error’, Mach. Learn. 52(3), 239–281.

Olson, C. A., Wu, N. C. & Sun, R. (2014), ‘A comprehensive biophysical description of pairwise epistasis throughout an entire protein domain’, Current Biology 24(22), 2643–2651.

Ortega, J. M. & Rheinboldt, W. C. (1970), Iterative Solution of Nonlinear Equations in Several Variables, SIAM.

Price, M. N., Wetmore, K. M., Waters, R. J., Callaghan, M., Ray, J., Liu, H., Kuehl, J. V., Melnyk, R. A., Lamson, J. S., Suh, Y., Carlson, H. K., Esquivel, Z., Sadeeshkumar, H., Chakraborty, R., Zane, G. M., Rubin, B. E., Wall, J. D., Visel, A., Bristow, J., Blow, M. J., Arkin, A. P. & Deutschbauer, A. M. (2018), ‘Mutant phenotypes for thousands of bacterial genes of unknown function’, Nature 557(7706), 503–509.

Riesselman, A. J., Ingraham, J. B. & Marks, D. S. (2018), ‘Deep generative models of genetic variation capture the effects of mutations’, Nature Methods 15(10), 816–822.

Robins, W. P., Faruque, S. M. & Mekalanos, J. J. (2013), ‘Coupling mutagenesis and parallel deep sequencing to probe essential residues in a genome or gene’, Proceedings of the National Academy of Sciences of the United States of America 110(9), E848–E857.

Romero, P. A., Krause, A. & Arnold, F. H. (2013), ‘Navigating the protein fitness landscape with Gaussian processes’, Proceedings of the National Academy of Sciences of the United States of America 110(3), E193–E201.

Romero, P. A., Tran, T. M. & Abate, A. R. (2015), ‘Dissecting enzyme function with microfluidic-based deep mutational scanning’, Proceedings of the National Academy of Sciences of the United States of America 112(23), 7159–7164.

Sali, A. & Blundell, T. L. (1993), ‘Comparative Protein Modelling by Satisfaction of Spatial Restraints’, Journal of Molecular Biology 234(3), 779–815.

Sauer-Eriksson, A. E., Kleywegt, G. J., Uhlén, M. & Jones, T. A. (1995), ‘Crystal structure of the C2 fragment of streptococcal protein G in complex with the Fc domain of human IgG’, Structure 3(3), 265–278.

Sloan, D. J. & Hellinga, H. W. (1999), ‘Dissection of the protein G B1 domain binding site for human IgG Fc fragment’, Protein Science 8(8), 1643–1648.

Song, H., Dai, R., Raskutti, G. & Barber, R. F. (2019), ‘Convex and non-convex approaches for statistical inference with noisy labels’, ArXiv e-prints.

Song, H. & Raskutti, G. (2018), ‘PUlasso: High-dimensional variable selection with presence-only data’, J. Am. Stat. Assoc. pp. 1–41.

Song, Y., Dimaio, F., Wang, R. Y. R., Kim, D., Miles, C., Brunette, T., Thompson, J. & Baker, D. (2013), ‘High-resolution comparative modeling with RosettaCM’, Structure 21(10), 1735–1742.

Suzek, B. E., Wang, Y., Huang, H., McGarvey, P. B. & Wu, C. H. (2015), ‘UniRef clusters: A comprehensive and scalable alternative for improving sequence similarity searches’, Bioinformatics 31(6).

Ward, G., Hastie, T., Barry, S., Elith, J. & Leathwick, J. R. (2009), ‘Presence-only data and the em algorithm’, Biometrics 65(2), 554–563.

Weile, J. & Roth, F. P. (2018), ‘Multiplexed assays of variant effects contribute to a growing geno-type–phenotype atlas’, Human Genetics 137(9), 665–678.

Weile, J., Sun, S., Cote, A. G., Knapp, J., Verby, M., Mellor, J. C., Wu, Y., Pons, C., Wong, C., Lieshout, N., Yang, F., Tasan, M., Tan, G., Yang, S., Fowler, D. M., Nussbaum, R., Bloom, J. D., Vidal, M., Hill, D. E., Aloy, P. & Roth, F. P. (2017), ‘A framework for exhaustively mapping functional missense variants’, Molecular Systems Biology 13(12), 957.

Wheeler, T. J. & Eddy, S. R. (2013), ‘Nhmmer: DNA homology search with profile HMMs’, Bioinformatics 29(19).

Wrenbeck, E. E., Bedewitz, M. A., Klesmith, J. R., Noshin, S., Barry, C. S. & Whitehead, T. A. (2019), ‘An Automated Data-Driven Pipeline for Improving Heterologous Enzyme Expression’, ACS Synthetic Biology 8(3), 474–481.

Wrenbeck, E. E., Faber, M. S. & Whitehead, T. A. (2017), ‘Deep sequencing methods for protein engineering and design’, Current Opinion in Structural Biology 45, 36–44.

Yang, K. K., Wu, Z. & Arnold, F. H. (2019), ‘Machine-learning-guided directed evolution for protein engineering’, Nature Methods 16(8), 687–694.

Yi, J., Hsieh, C.-J., Varshney, K. R., Zhang, L. & Li, Y. (2017), Scalable Demand-Aware recommendation, in I. Guyon, U. V. Luxburg, S. Bengio, H. Wallach, R. Fergus, S. Vishwanathan & R. Garnett, eds, ‘Advances in Neural Information Processing Systems 30’, Curran Associates, Inc., pp. 2412–2421.

## References

[1] Lung-Fei Lee. Specification error in multinomial logit models: Analysis of the omitted variable bias. J. Econom., 20(2):197–209, November 1982.

